# DTCAMD, a disease-associated truncated form of CRYAB in neovascular age-related macular degeneration: a potential driver of subretinal fibrosis

**DOI:** 10.64898/2026.10.06.756917

**Authors:** Eunhye Yu, Sun Mi Gu, Jonghwa Jin, Jung Kee Min, Jaesuk Yun

## Abstract

**Purpose:** Subretinal fibrosis causes irreversible vision loss in neovascular age-related macular degeneration (nAMD) and remains poorly controlled by anti–vascular endothelial growth factor (VEGF) therapy. We investigated the role of αB-crystallin (CRYAB) in angiogenic and fibrotic pathology and evaluated disease-associated CRYAB truncation as a therapeutic target.

**Methods:** Aqueous humor from patients with nAMD was analyzed by quantitative proteomics and receiver operating characteristic (ROC) analysis. CRYAB function was assessed using knockout and knockdown approaches in a choroidal neovascularization (CNV) mouse model. CRYAB truncation and associated proteins were examined by immunoblotting, co-immunoprecipitation, and liquid chromatography–tandem mass spectrometry (LC–MS/MS). C-terminal truncation was further examined in mouse retinal tissue using antibodies generated against full-length CRYAB and its C-terminal region. Primary human retinal pigment epithelial (hRPE)-THP-1 co-cultures were used to examine CRYAB truncation under fibrosis-associated conditions. Pharmacological effects were tested using (±)-threo-3-methylglutamic acid (T3MG) and 4-(2-aminoethyl)benzenesulfonyl fluoride hydrochloride (AEBSF).

**Results:** CRYAB was upregulated in the aqueous humor of patients with nAMD. Higher CRYAB levels were associated with persistent lesions after initial anti-VEGF treatment. CRYAB knockout or knockdown reduced CNV pathology and decreased VEGFA and α-smooth muscle actin (αSMA) expression, without a significant substrain effect between C57BL/6N and C57BL/6J mice. A Ser45-phosphorylated truncated CRYAB species was detected in mouse laser-induced CNV model. We termed this species disease-associated truncated CRYAB in AMD (DTCAMD). Peptide mapping showed reproducible loss of distal C-terminal peptide coverage in truncated CRYAB. Consistent with this finding, C-terminal antibody recognition was reduced in CNV lesions. In hRPE–THP-1 co-cultures, DTCAMD accumulated with increased αSMA expression. Proteomic analysis identified Prss1 as a candidate CRYAB-associated serine protease linked to truncation under CNV conditions. T3MG and AEBSF reduced DTCAMD-associated signals and CNV pathology.

**Conclusions:** CRYAB contributes to angiogenic and fibrotic pathology in CNV. Disease-associated CRYAB truncation, particularly Ser45-phosphorylated DTCAMD, represents a potential therapeutic target for fibrotic remodeling in nAMD.

## Introduction

Neovascular age-related macular degeneration (nAMD) is characterized by abnormal choroidal neovascular growth into the subretinal space, leading to fluid exudation, hemorrhage, and progressive loss of central vision^1^. Intravitreal anti– vascular endothelial growth factor (VEGF) therapy effectively suppresses pathological angiogenesis and has substantially improved visual outcomes^2–6^. Its effect on subretinal fibrosis, however, remains limited. Approximately 45% of patients develop fibrotic scarring within two years of anti-VEGF treatment highlighting a persistent therapeutic gap and the need to define mechanisms that drive scar formation independently of VEGF-mediated angiogenesis ^7,8^.

Proteomic profiling of aqueous humor from patients with nAMD revealed increased levels of αB-crystallin (CRYAB), a small heat shock protein with well-established cytoprotective and neuroprotective functions^9^. In parallel, immunoblotting in experimental nAMD models identified a prominent truncated CRYAB species that increased together with full-length CRYAB. CRYAB normally functions as a molecular chaperone that limits protein misfolding and aggregation^10–14^. The appearance of a truncated species under disease conditions suggests that this protective function may be altered within the retinal microenvironment. Although the protease responsible for this processing has not been established, the serine protease PRSS1/Prss1 was increased in the same experimental context, suggesting a potential role in disease-associated CRYAB processing.

A possible basis for this functional shift lies in disruption of the CRYAB C-terminal domain (CTD). The CTD contains the conserved IXI/V motif (Ile159–Pro160–Ile161), which engages the hydrophobic β4–β8 groove within the central α-crystallin domain (ACD) and contributes to the regulation of intermolecular interactions and oligomeric stability^15,16^ (Fig. S1A-C). Disruption of this CTD–ACD interaction can expose hydrophobic surfaces and alter protein–protein interactions. Structural perturbations of CRYAB, including the D109A mutation and stress-associated truncation, have likewise been associated with impaired chaperone activity and increased aggregation propensity^17,18^. In our peptide-mapping analysis, the truncated CRYAB species showed a truncation pattern consistent with a boundary near the Arg149–Lys150 region. Such a truncation would remove the downstream C-terminal segment containing the IXI/V motif.

Based on these observations, we defined this processed species as disease-associated truncated CRYAB in AMD (DTCAMD). DTCAMD was associated with Ser45 phosphorylation, suggesting that truncation and phosphorylation may occur within the same disease-associated CRYAB population. Loss of the C-terminal regulatory region could increase accessibility of the β4–β8 hydrophobic groove and alter the interaction profile of CRYAB. We therefore examined whether Ser45-phosphorylated DTCAMD is associated with fibrosis-related proteins, including α-smooth muscle actin (αSMA), and whether its accumulation tracks with fibrotic remodeling under CNV conditions.

In this study, we investigated the presence and pathological relevance of DTCAMD using complementary experimental approaches. We examined CRYAB-associated proteins, mapped the molecular features of the truncation, and assessed its relationship with Ser45 phosphorylation and fibrotic markers. We also tested whether genetic or pharmacological suppression of the CRYAB axis reduces DTCAMD accumulation and attenuates VEGFA and αSMA responses. These experiments were designed to determine whether disease-associated CRYAB processing contributes to CNV-associated fibrotic pathology and whether this pathway represents a therapeutic target beyond VEGF inhibition.

## Materials and methods

### 1. Ethical approval and patient consent

The clinical study protocol was approved by the Institutional Human Experimentation Committee Review Board of Ulsan University Hospital, Ulsan, Republic of Korea (NON2020-004 and UUH2018-03-006). All procedures were conducted in strict accordance with the ethical standards laid down in the 1964 Declaration of Helsinki and its later amendments. Informed consent was obtained from all patients prior to their inclusion in the study.

### 2. Animals and ethical approval

All animal experiments were performed in accordance with the Association for Research in Vision and Ophthalmology (ARVO) Statement for the Use of Animals in Ophthalmic and Vision Research and followed the ARRIVE guidelines (https://arriveguidelines.org/). The study protocol was approved by the Animal Experimentation Ethics Committee of Chungbuk National University, Cheongju, Republic of Korea (CBNUA-2069-23-02).

Male WT C57BL/6N and CRYAB knockout (KO) mice were maintained under standard housing conditions with a 12-h light/dark cycle and randomly assigned to cages (4–5 animals per cage; n = 5–8 per group). To generate CRYAB KO mice, transgenic mice were bred with C57BL/6N mice (Daehan BioLink, Chungbuk, Korea) to obtain heterozygous F1 offspring. These were subsequently bred to produce F2 CRYAB KO mice, which were genotyped using tail DNA at 3–4 weeks of age. Only male CRYAB KO mice were used for experiments, and F2 inbred mice were further bred to obtain the F3 generation.

### 3. Patient diagnosis and aqueous humor sampling

All participants underwent a complete ophthalmic examination that included the best-corrected visual acuity assessment, fundus examination using slit-lamp ophthalmoscopy, fundus photography (Daytona, Optos, Inc., Marlborough, MA, USA), swept-source OCT, and OCT angiography (OCTA) (DRI OCT Triton; Topcon Corporation, Tokyo, Japan), and the nAMD diagnosis was given when CNV was detected via fluorescein angiography (FA) and indocyanine green angiography (ICGA) (Heidelberg Retina Angiograph Spectralis, Heidelberg Engineering, Heidelberg, Germany). nAMD was diagnosed by fundus examination, including one or more features such as subretinal fluid (SRF) or intraretinal fluid (IRF), subretinal hemorrhage, hyper-reflective lesions, and retinal pigmented epithelial detachment (PED) in the macular area, using fundus photography and OCT. A total of 62 participants were enrolled in this study, with 48 patients in the nAMD group and 14 controls without nAMD. The control group consisted of patients without nAMD who were randomly selected from those undergoing macular hole surgery. The selection was based on age, similar to that of the nAMD group. All 48 patients with nAMD received an anti-VEGF agent (aflibercept) intravitreal injection treatment three times at monthly intervals; all eye examinations except FA/ICGA were repeated 1 week after the injection and 1 month after the third injection.

Aqueous humor samples were collected at the beginning of the surgery or intravitreal injection procedure using the following procedure: after sterile surgical draping was performed, a sterile eyelid speculum was placed, and 1–2 drops of 0.5% proparacaine hydrochloride (Alcaine, Alcon, Ft. Worth, TX, USA) and 5% povidone iodine were instilled immediately before surgery or injection. Anterior chamber paracentesis was performed using a 30-gauge needle mounted on a 1-mL syringe through a puncture at the peripheral cornea under a surgical microscope. Aqueous humor sample volumes ranged from 50 to 100 µL. The samples were transferred to microtubes and immediately stored at –70 °C.

### 4. Measurement of clinical parameters

CMT was measured using OCT; measurement at the initial visit was defined as pre-CMT, and measurement 1 month after three injections was defined as post-CMT. When a PED was present at the fovea, CMT, including PED from the outer segment/RPE layer to the internal limiting membrane, was defined as CMT (including PED). The SRF area was also measured using OCT, and the measurement at the time of diagnosis was defined as the pre-SRF area. Lesion resolution was evaluated based on an OCT exam 1 month after three injections. The CNV area was measured using ICGA and OCTA at the initial visit. ICGA images in which the CNV was clearly visible were selected, and the inner border of the CNV was manually outlined and measured using the area-measuring function of the built-in software. The CNV shown in the choriocapillaris layer in the OCTA image obtained on the same day was also measured using the software embedded in the OCT instrument. The average CNV area measured using the two devices was defined as the CNV area^19^. In addition, the presence or absence of OCT biomarkers (SRF, IRF, subretinal hemorrhage, hyperreflective lesions, and PED) visible on OCT in patients with nAMD was used as a criterion for determining treatment effectiveness.

### 5. Proteomic analysis of aqueous humor

The analysis of aqueous humor samples, including protein identification and quantification, was conducted as described in our previous study ^20^. Briefly, samples were obtained from 28 participants (14 patients with nAMD and 14 controls) following ethical guidelines and institutional approvals (NON2020-004 and UUH2018-03-006). LC-MS/MS was employed using TMT-based quantitative proteomics. For the current study, the focus was extended to include CRYAB, which was identified in the previous dataset but not specifically analyzed.

### 6. Quantification of CRYAB by ELISA

The concentration of CRYAB in the aqueous humor was measured using a commercially available enzyme-linked immunosorbent assay kit (#ADI-960-074, Enzo Life Sciences, Inc., Farmingdale, NY, USA) according to the manufacturer’s instructions. All samples (34 patients with nAMD) were analyzed in duplicate. The levels were determined with a microplate reader (VersaMax Microplate Reader, Molecular Devices, Orleans, CA, USA).

### 7. Genotyping of the Crb1^rd^^8^ mutation by PCR

The presence of the Crb1^rd^^8^ mutation in the C57BL/6N background was assessed by PCR (Fig. S2A). Because C57BL/6 substrains may differ in genetic background, we additionally performed confirmatory experiments in C57BL/6J mice to assess the robustness of CNV/fibrosis phenotypes and the response to CRYAB inhibition across substrains. DNA was isolated from mouse tail biopsies using the REDExtract-N-Amp™

Tissue PCR Kit (Sigma-Aldrich) following the manufacturer’s instructions. To differentiate between the wild-type (WT) and *rd8* mutant alleles, PCR amplification was performed using primers previously described by Mehalow et al^21^. The primer sequences were as follows: m*Crb1* mF1: GTGAAGACAGCTACAGTTCTGATC, m*Crb1* mF2: GCCCCTGTTTGCATGGAGGAAACTTGGAAGACAGCTACAGTTCTTCTG, m*Crb1* mR: GCCCCATTTGCACACTGATGAC. Each PCR reaction was carried out in a 10 µL volume containing approximately 2 µL of Tissue Extract, 5 µL of REDExtract-N-Amp™ PCR Reaction Mix (Sigma-Aldrich), and primers at the following concentrations: 1.6 μM each of forward and reverse primer for WT allele, and 0.8 μM of forward and 1.6 μM of reverse primer for *rd8* mutant allele. Reactions initially were denatured at 94°C for 5 minutes followed by 35 cycles at 94°C for 30 seconds, 65°C for 30 seconds, 72°C for 30 seconds and a final extension at 72°C for 7 minutes. Amplicons were separated on a 2% agarose gel and visualized under UV light. The expected amplicon sizes were 220 bp for the WT allele and 244 bp for the *rd8* allele.

### 8. *In vivo* electroporation for CRYAB knockdown

To induce CRYAB knockdown in the mouse retina, *in vivo* electroporation was performed using the NEPA21 Super Electroporator (Nepagene, Chiba, Japan) (Fig. S2B-C). A CRYAB-specific siRNA pool containing two duplexes (12955-1 and 12955-2; Bioneer, Daejeon, Republic of Korea) mixed at a 1:1 ratio was used. For 12955-1, the oligonucleotide sequences were 5′-AGAAGGACAGAUUCUCUGU-3′ and 5′-ACAGAGAAUCUGUCCUUCU-3′; for 12955-2, the sequences were 5′-GACUAGUGCUGAAGCUUAU-3′ and 5′-AUAAGCUUCAGCACUAGUC-3′. All strands contained 3′-dTdT overhangs. First, CRYAB-specific siRNA or scrambled control siRNA (Bioneer, Daejeon, Republic of Korea) was prepared at a concentration of 1 µg/µL and intravitreally injected into the left eye of each mouse. Immediately following the injection, electric pulses were delivered using 10-mm diameter tweezer electrodes (CUY650P10; Nepagene). To facilitate the delivery of siRNA into the RPE, the negative electrode (cathode) was positioned on the left eye (injection site), and the positive electrode (anode) was placed on the contralateral right eye^22^. The electroporation parameters were set as follows: voltage, 80 V; pulse length, 50 ms; pulse interval, 950 ms; and number of pulses, 5. The efficacy of CRYAB knockdown was evaluated at 7-and 14-days post-transfection (Fig. S2D-E).

### 9. Laser-Induced CNV model and drug administration

For all procedures, animals were anesthetized via intraperitoneal injection of 1.2% avertin. The pupils were dilated using a combination of 0.5% tropicamide and 0.5% phenylephrine (Mydrin-P; Santen, Osaka, Japan). Laser photocoagulation was performed on both eyes of each animal at the retina, with a distance of 2–3 optic disc diameters surrounding the optic nerve, avoiding blood vessels. Laser spots were created with a spot diameter of 45 µm using the 78 diopters lens (VOLK, Mentor, OH, USA), 300 mW power, and 100 ms pulse duration using a laser indirect ophthalmoscope (OcuLight®GL; Green 532 nm Laser; IRIDEX, Mountain View, CA, USA).

T3MG, which can block the interaction between CRYAB and VEGFA^23^, was obtained from Tocris Bioscience (0811, Bristol, UK). AEBSF, which is an irreversible serine protease inhibitor, was purchased from Sigma-Aldrich (A8456, St. Louis, MO, USA). The compounds were dissolved in PBS. Aflibercept (Eylea; Bayer AG, Berlin, Germany), an anti-VEGF agent, was used as a positive control. Before intravitreal injection, a small amount of vitreous fluid was removed after puncture using a 30-gauge needle, being careful not to touch the lens on the limbus. Then, mice were intravitreally injected at the same puncture site using a 32-gauge needle attached to a 5 µL Hamilton micro syringe (Hamilton, Reno, NV, USA) containing: 1 µL PBS or T3MG (25 µg in 1 µL PBS), AEBSF (25 µg in 1 µL PBS), T3MG + AEBSF (25 µg in 0.5 µL PBS + 25 µg in 0.5 µL PBS), or aflibercept (40 µg/1 µL).

### 10. Evaluation of CNV and fibrotic lesions

The experimental mice were tested after 1 week. For all procedures, the animals were anesthetized via an intraperitoneal injection of 1.2% avertin. The pupils were dilated using a combination of 0.5% tropicamide and 0.5% phenylephrine (Mydrin-P). Following the intraperitoneal injection of 2.5% fluorescein sodium (0.5 mL) (Novartis, Basel, Switzerland), retinal imaging was performed using a UNIENDO fundus camera (Conecson, Ulsan, Korea). After administration, retinal images were recorded in video mode to capture fluorescence leakage, and evaluation was performed 1 min later to assess the area of leakage around the new chorioretinal vessels.

To further visualize and quantify the CNV area, choroidal flat mounts were prepared. Enucleated eyes were fixed in 4% paraformaldehyde for 1 h, and the RPE/choroid complexes were dissected. After blocking with 5% bovine serum albumin (BSA) containing 0.5% Triton X-100 for 1 h, the tissues were incubated overnight at 4°C with isolectin B4 (IB4) conjugated to Alexa Fluor 594 (1:100; Invitrogen, Waltham, MA, USA). The flat mounts were mounted and imaged using a Zeiss Axio Imager A2 microscope (Carl Zeiss, Oberkochen, Germany). The CNV area was quantified using ImageJ software.

Laser photocoagulation spots were identified on digital red-free retinal images using spectral-domain OCT (Eyemera OCT; IISCIENCE, CA, USA). Retinal OCT scans were performed twice in the vertical and horizontal planes based on the center of the laser spot. Hyper-reflective lesions were confirmed on OCT images scanned in the vertical and horizontal planes. The inner border of the lesion was manually outlined, and the cross-sectional area of hyper-reflective lesions was measured using ImageJ software (version 1.53, https://imagej.nih.gov/ij/index.html).

### 11. Protein extraction

Mouse retinal, RPE, and choroidal tissues were isolated via sonication in lysis buffer (iNtRON Biotechnology, Seongnam, Korea) and centrifuged at 13,000 rpm for 25 min at 4 °C. The resulting supernatant was used as the total protein extract. Cells from hRPE–THP-1 co-cultures were harvested and lysed in lysis buffer (iNtRON Biotechnology, Seongnam, Korea), followed by centrifugation at 13,000 rpm for 25 min at 4 °C. The resulting supernatants were used as cell lysates for immunoblot analysis.

### 12. Immunoblot analysis

Tissue or cell lysates were resolved using 12% glycine SDS-PAGE gel and transferred onto a polyvinylidene difluoride membrane, which was incubated overnight at 4 °C with αSMA (Abcam, Cambridge, UK; ab5694, 1:1,000), VEGFA (Abcam, ab46154, 1:1,000), CRYAB (Abcam, ab13496 and ab13497, 1:2,500), p-CRYAB (Ser45 and Ser59; Invitrogen, PA1-011 and PA1-012, 1:2,500), and GAPDH (Cell Signaling Technology, MA, USA; 2118S, 1:5,000) antibodies. The blots were then incubated with the corresponding conjugated goat anti-rabbit (Cell Signaling Technology, 7074S, 1:5,000) or horse anti-mouse IgG-horseradish peroxidase (HRP) (Cell Signaling Technology, 7076S, 1:5,000) secondary antibodies. Immunoreactive proteins were detected using an ECL substrate (Millipore, Burlington, MA, USA) and visualized using a FUSION Solo S chemiluminescence detection system (Vilber Lourmat, Collegien, France).

### 13. LC–MS/MS peptide mapping of CRYAB

For CRYAB peptide mapping, mouse eye lysates from naïve and laser-induced choroidal neovascularization (CNV) eyes were separated by SDS-PAGE and stained with InstantBlue™ Coomassie Protein Stain (Abcam, ab119211). Gel regions corresponding to full-length CRYAB from naïve eyes, full-length CRYAB from laser-treated eyes, and truncated CRYAB from laser-treated eyes were independently excised and subjected to in-gel digestion followed by LC–MS/MS analysis. Peptide analysis was performed using liquid chromatography coupled to a hybrid Fourier transform (FT) Orbitrap mass spectrometer. MS/MS data were analyzed using the Mascot server software (Matrix Science, London, UK) against the CRYAB protein sequence. Trypsin was specified as the proteolytic enzyme, with carbamidomethylation of cysteine as a fixed modification and oxidation of methionine as a variable modification. Identified peptides were mapped to the CRYAB amino acid sequence to compare peptide coverage among the analyzed gel regions.

### 14. Co-Immunoprecipitation (Co-IP) assay

Co-IP assays were performed using the Pierce Classic Magnetic IP/Co-IP Kit (Thermo Fisher Scientific) according to the manufacturer’s instructions. Briefly, mouse eyes lysate containing 500 µg total protein (quantified using the Bradford assay) was incubated with 2 µg CRYAB or Prss1 antibodies for IP overnight at 4 °C. Thereafter, the antigen/antibody complex was incubated with 25 µL Pierce Protein A/G magnetic beads for 1 h at room temperature. The beads were then washed twice with IP lysis/wash buffer and once with distilled water and heated at 100 °C for 5 min to elute the antigen/antibody complexes from the beads, followed by western blotting.

### 15. LC–MS/MS identification of CRYAB-associated proteins

To identify proteins associated with CRYAB, mouse eye lysates were subjected to CRYAB Co-IP. The recovered protein samples were mixed with 5× sample buffer, heated at 100 °C for 1 min, and separated by SDS-PAGE. Electrophoresis was performed at 130 V until the proteins migrated approximately 1 cm into the resolving gel, and the protein-containing gel region was excised and subjected to in-gel digestion. The resulting peptide samples were purified and analyzed using an EASY-nLC 1000 system coupled to a Q-Exactive Hybrid Quadrupole Orbitrap mass spectrometer (Thermo Fisher Scientific). Peptides were separated by nano-LC using an acetonitrile gradient at a flow rate of 300 nL/min over a total analysis time of 70 min. Protein identification was performed against a Mus musculus database using Proteome Discoverer 2.2 software.

### 16. Immunofluorescence staining

Mice were euthanized 7 days after laser photocoagulation. Enucleated eyes from mice were fixed in 4% paraformaldehyde (PFA) for 2 h at 4 °C, followed by cryoprotection in 30% sucrose for at least 12 h at 4 °C. The tissues were then embedded in optimal cutting temperature (OCT) compound, frozen, and sectioned at a thickness of 15 µm using a cryostat (CM1850; Leica Microsystems, Wetzlar, Germany). Sections were permeabilized with PBS containing 0.3% Triton X-100 and 0.3% hydrogen peroxide and blocked with 5% BSA containing 1% goat serum for 1 h at room temperature. The sections were incubated overnight at 4 °C with antibodies against CRYAB (Abcam, ab13496 and ab13497, 1:200) diluted in 5% BSA. After three washes with PBS, the sections were incubated for 2 h at room temperature with Alexa Fluor 488-conjugated anti-mouse IgG and Alexa Fluor 568-conjugated anti-rabbit IgG secondary antibodies diluted in 5% BSA. Nuclei were counterstained with DAPI (300 nM; Sigma-Aldrich) for 20 min. The sections were mounted on glass slides (Matsunami, Osaka, Japan) using mounting medium (Vector Laboratories, Inc., San Francisco, CA, USA), and images were acquired using a confocal microscope (LSM 880; ZEISS, Oberkochen, Germany).

### 17. hRPE and THP-1 co-culture

To study disease-associated CRYAB processing in a more physiologically relevant setting, a co-culture model was established using hRPE cells (RPE001; Neuromics, MN, USA) and THP-1 cells. hRPE cells were cultured in human epithelial cell medium (ECM001; Neuromics) supplemented with 1% penicillin-streptomycin. THP-1 cells were differentiated into macrophage-like cells by treating them with 200 ng/mL PMA overnight, followed by 1 μg/mL LPS treatment for 24 h. After differentiation, THP-1 cells were co-cultured with hRPE cells at a 1:2 ratio in 12-well plates. Co-cultures were maintained for 3, 24, 48, and 72 h to assess time-dependent changes. CRYAB, phosphorylated CRYAB, and truncated CRYAB species were subsequently analyzed by western blotting to evaluate time-dependent CRYAB processing under inflammatory co-culture conditions.

### 18. Statistical analysis

The ROC curve was plotted using IBM SPSS Statistics for Windows (version 21.0; IBM Corp, Armonk, NY, USA) and shown as mean ± standard deviation. All other results are shown as mean ± SEM. Data distribution within all separate groups were validated by the Shapiro–Wilk normality test. An unpaired two-tailed t-test was used for comparisons between two groups. If the normality assumption was not met, the non-parametric Mann–Whitney test was used. One-Way ANOVA followed by Holm– Šídák’s multiple-comparison test was used for comparison of more than two groups, if normality test was not passed the non-parametric Kruskal–Wallis test was performed. Two-way or three-way ANOVA followed by Holm–Šídák’s multiple-comparisons test was used, as appropriate, to evaluate the effects and interactions of multiple experimental factors. The statistical significance level was set at 0.05, unless otherwise stated. Statistical analyses were performed with SigmaPlot 14 software (Systat Software, San Jose, CA, USA) or GraphPad Prism (GraphPad Software, Boston, MA, USA).

## Results

### 1. Differential expression of CRYAB in the aqueous humor of patients with nAMD

Proteomic analysis of the aqueous humor samples revealed significant differences in protein expression profiles between patients with nAMD and controls. As part of this study, our previously identified datasets^20^ were reanalyzed with a focus on CRYAB and its potential association with angiogenic and fibrotic pathology. Using tandem mass tag (TMT, Thermo Fisher Scientific, Waltham, MA, USA)-labeled quantification across three technical replicates, we identified a total of 127 differentially expressed proteins meeting the following criteria: t-test *p*-value < 0.05, confidence level > 95%, a minimum of two unique peptides, FDR < 5%, and fold-change thresholds of log_2_ > 1.5 or < -1.5. Among these proteins, CRYAB was notably upregulated in the nAMD group, exhibiting a significant fold-change compared to controls. Volcano plot analysis further highlighted CRYAB as one of the most differentially expressed proteins (Fig. 1A,B). These data identified CRYAB as a prominently increased protein in the aqueous humor of patients with nAMD and prompted further investigation of its pathological relevance.

**Figure 1.**
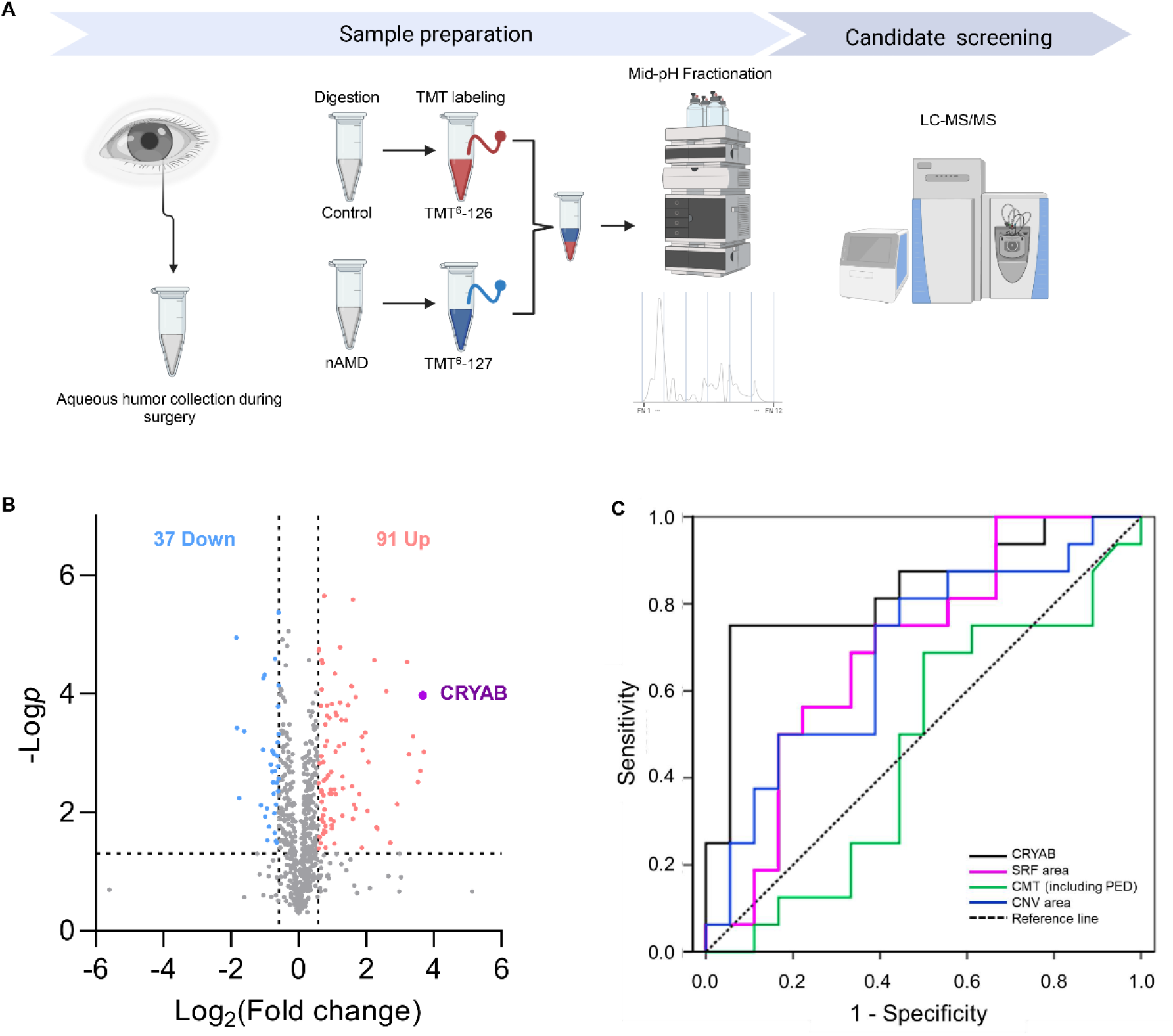
CRYAB as a predictive marker for nAMD treatment outcomes in human aqueous humor. (A) Schematic overview of the clinical study design and analysis workflow. The illustration was created in BioRender. Yu, E. (https://BioRender.com/p5mc14x). (B) Differential expression of CRYAB in the aqueous humor of patients with nAMD and controls. Quantitative proteomics analysis identified CRYAB as significantly upregulated in patients with nAMD compared to controls. The cutoff range of protein quantitation is as follows: t-test, p value < 0.05, minimum confidence level > 95%, more than 2 unique peptides, FDR < 5%, and fold-change thresholds (expressed as log2 ratio) of > 1.5 or < -1.5. Data were analyzed using Perseus (version 1.5.8.2) for the analysis of differentially expressed proteins and statistical analysis. The cutoff range for significant fold change (FC) and T-test p-values were set as ±1.5 and 0.05, respectively. (C) Comparison of ROC curves using CRYAB concentration in aqueous humor (black line), SRF area (magenta line), central macular thickness (CMT) (including PED) (green line), and CNV area (blue line) before treatment to predict resolution or persistence of subretinal and intraretinal lesions (hyper-reflective lesions, SRF, and PED) after initial three intravitreal anti-VEGF agent (aflibercept) injections after nAMD diagnosis. The area under the curve values were 0.830 for CRYAB, 0.694 for SRF area, 0.688 for CNV area, and 0.467 for CMT (including PED). CRYAB was statistically significant (p = 0.001), but the others (SRF area, CNV area, and CMT) did not show statistically significant values (SRF area: p = 0.053, CNV area: p = 0.062, CMT: p = 0.743).

### 2. CRYAB levels in aqueous humor outperformed traditional markers in predicting lesion resolution in patients with nAMD treated with anti-VEGF injection therapy

This study included 34 patients diagnosed with nAMD who were treated with an anti-VEGF agent (aflibercept) via intravitreal injections (Table 1). Following the initial three injections, optical coherence tomography (OCT) showed complete lesion resolution in 18 patients, whereas 16 patients had persistent lesions, including residual subretinal fluid (SRF), hyper-reflective lesions, or pigment epithelial detachment (PED) (Table 2). ROC curves were used to predict the resolution or persistence of subretinal and intraretinal lesions (hyperreflective lesions, SRF, and PED) after the three initial intravitreal injections. The ROC curves were used to compare the predictive abilities of CRYAB levels in the aqueous humor with those of several typical markers of nAMD (initial SRF area, CNV area, and central macular thickness [CMT], including PED). CRYAB demonstrated superior sensitivity and specificity compared to the other methods. The areas under the curve were 0.830 for CRYAB, 0.694 for SRF, 0.688 for CNV, and 0.467 for CMT (including PED). CRYAB expression was statistically significant (*p* = 0.001), but the other markers (SRF area, CNV area, and CMT, including PED) did not show statistically significant differences (SRF area, *p* = 0.053; CNV area, *p* = 0.062; CMT, *p* = 0.743) (Fig. 1C). Within this cohort, aqueous humor CRYAB showed greater discriminatory performance than the OCT-derived parameters examined.

**Table 1.** Demographic and clinical characteristics of patients with nAMD. Data are presented as mean ± standard deviation. BCVA, best-corrected visual acuity; CNV, choroidal neovascularization; CMT, central macular thickness; CRYAB, crystallin alpha B; logMAR, logarithm of the minimal angle of resolution; PED, pigmented epithelial detachment; SRF, subretinal fluid.

| Characteristics | Eyes with nAMD (n = 34) |
| --- | --- |
| Age (years) | 66.44 ± 9.38 |
| Men/women | 10/24 |
| CRYAB (ng/mL) | 51.25 ± 53.06 |
| Pre-injection BCVA (logMAR) | 0.50 ± 0.37 |
| Post-injection BCVA (logMAR) | 0.34 ± 0.28 |
| Pre-CMT (including PED) (μm) | 370.59 ± 119.12 |
| Pre-SRF area (mm<sup>2</sup>) | 7.53 ± 4.98 |
| Pre-CNV area (mm<sup>2</sup>) | 3.95 ± 3.56 |
| Post-CMT (including PED) (μm) | 205.88 ± 56.45 |

**Table 2.**
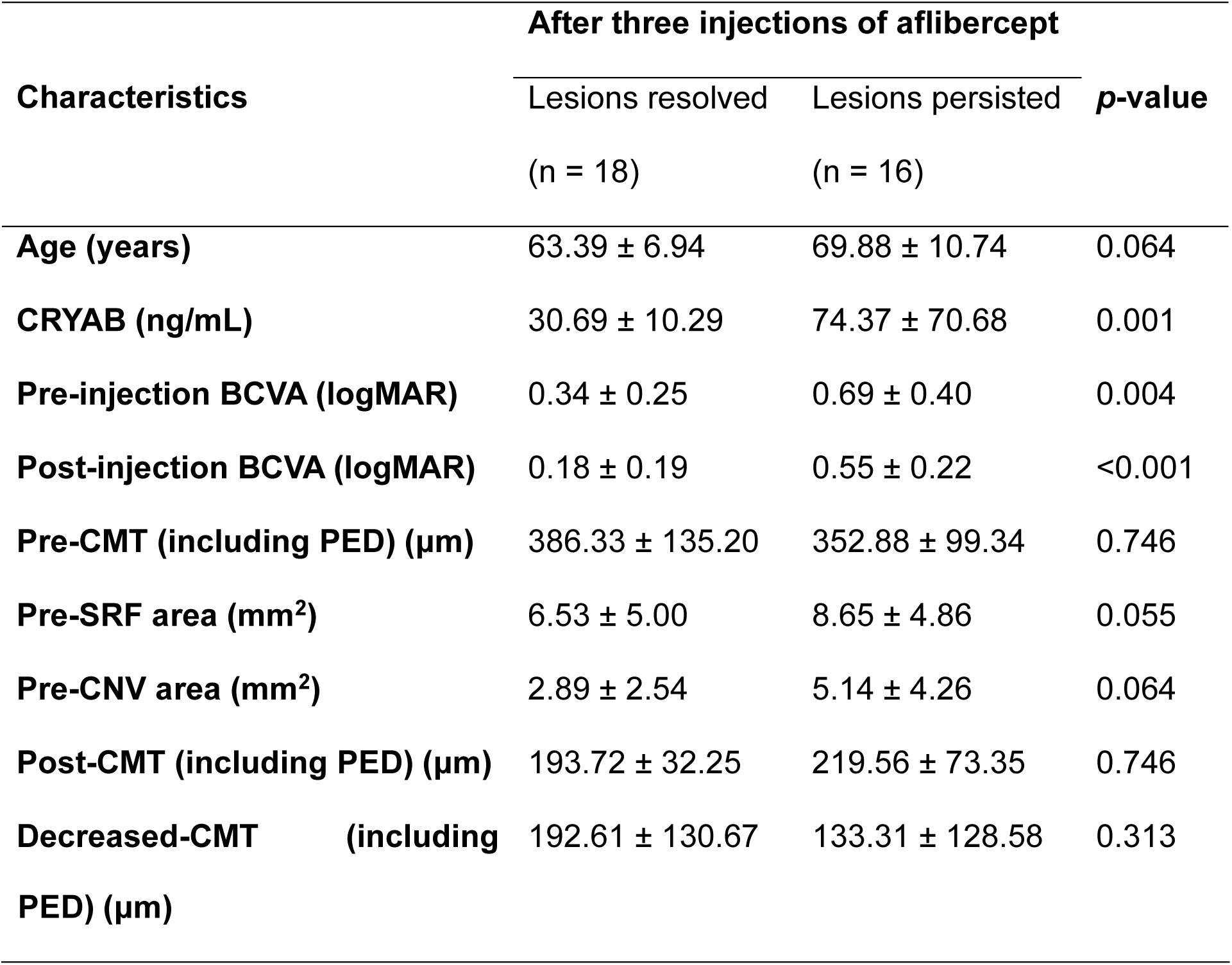
Comparison of age, CRYAB levels in aqueous humor, visual acuity, CMT, SRF area, CNV area, and CMT change between subretinal and resolved and persisting intraretinal lesion (hyper-reflective lesions, SRF, and PED) groups. Data are presented as mean ± standard deviation. p-values were derived using Mann–Whitney U test. BCVA, best-corrected visual acuity; CNV, choroidal neovascularization; CMT, central macular thickness; CRYAB, αB-crystallin; logMAR, logarithm of the minimal angle of resolution; PED, pigmented epithelial detachment; SRF, subretinal fluid.

### 3. Genetic ablation and knockdown of CRYAB attenuate CNV and fibrotic pathology *in vivo*

To determine whether CRYAB contributes functionally to CNV-associated pathology, we employed a laser-induced CNV model in CRYAB knockout (KO) mice (Fig. 2A). Compared with wild-type (WT) controls, CRYAB-deficient mice exhibited a marked reduction in CNV lesion size and vascular leakage, as assessed by fluorescein angiography (Fig. 2B). Consistently, OCT imaging revealed a significant decrease in hyper-reflective lesions in CRYAB KO eyes (Fig. 2C). At the molecular level, western blot analysis demonstrated significantly lower expression of αSMA and VEGFA in CRYAB KO mice than in WT controls (Fig.2D-F).

**Figure 2.**
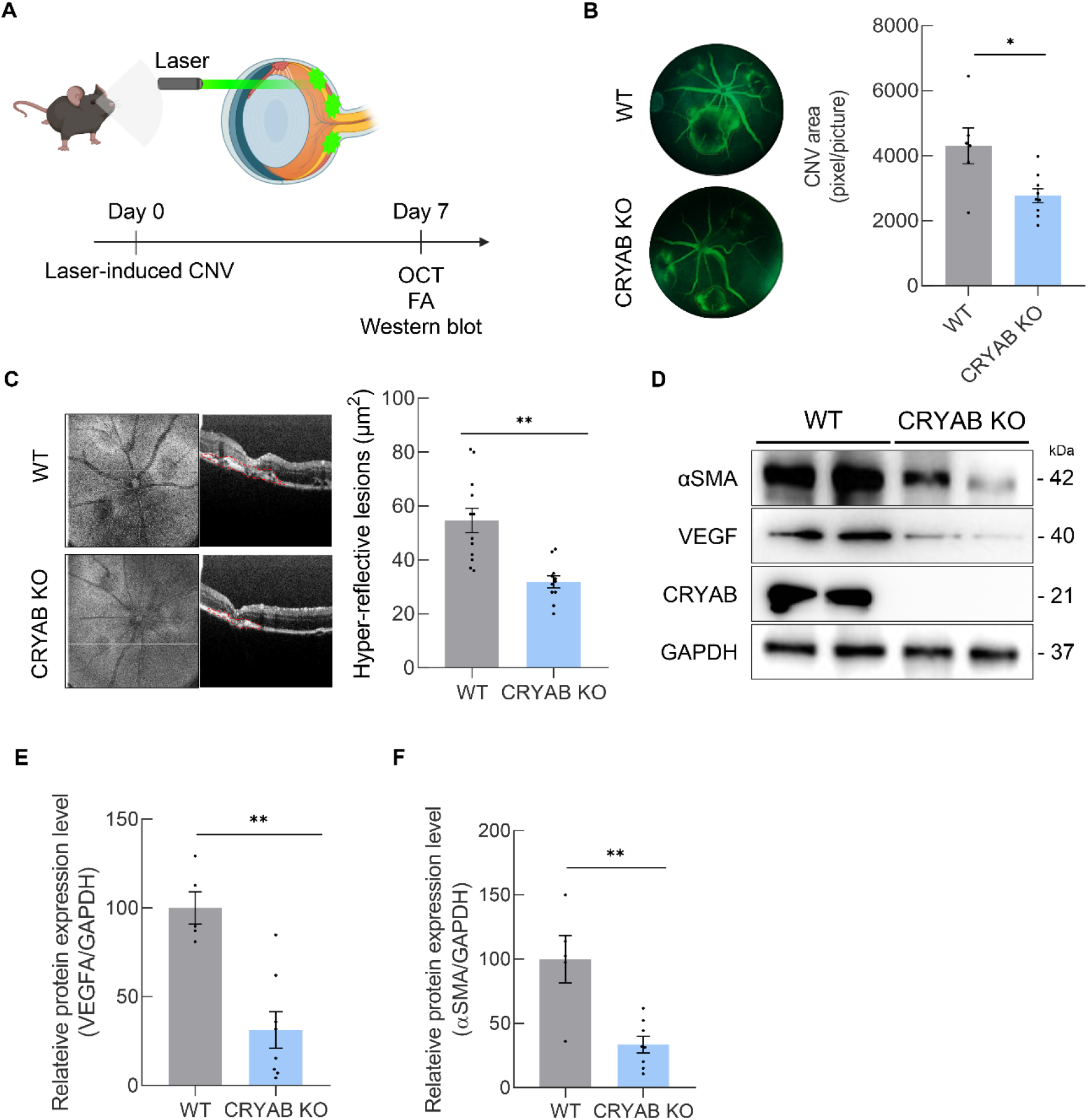
CRYAB deficiency attenuates CNV-associated angiogenic and fibrotic pathology *in vivo*. (A) Schematic illustration of CNV induction and analysis. (B) Representative fluorescein angiography (FA) images of the WT and CRYAB KO mouse eyes 1 week after laser-induced CNV. Bar graph showing fluorescence leakage size measured from FA images (n = 6–9). (C) Representative optical coherence tomography (OCT) images of WT and CRYAB KO mouse eyes 1 week after laser-induced CNV. Bar graph showing hyper-reflective lesions measured from OCT scan (n = 11–12). (D,E) Representative western blot showing αSMA and VEGFA proteins in whole-tissue lysates of laser-induced mouse eye extracts 1 week after laser exposure. GAPDH was used as loading control (n = 5-8). Data are presented as mean ± SEM and were analyzed using unpaired two-tailed Student’s t-tests. *p < 0.05 and **p < 0.01 vs. WT mice.

Because the CRYAB KO mice were maintained on a C57BL/6N background, which can carry the Crb1^rd^^8^ mutation, we next examined whether substrain-related retinal differences could influence the observed phenotype. Laser injury induced comparable CNV lesion size and subretinal hyper-reflective areas in C57BL/6N and C57BL/6J mice, consistent with previous reports indicating that laser-induced CNV development is not markedly affected by C57BL/6 substrain background^24^. CRYAB knockdown significantly reduced both SHRM and CNV area in C57BL/6N and C57BL/6J mice (Fig 3A-E). Consistently, western blot analysis showed reduced VEGFA and αSMA expression following CRYAB knockdown in both substrains (Fig. 3F-I). No significant substrain main effect or substrain-related interaction was observed for these structural or molecular outcomes. The effects of CRYAB suppression were therefore reproducible in both C57BL/6N and C57BL/6J mice under the experimental conditions tested.

**Figure 3.**
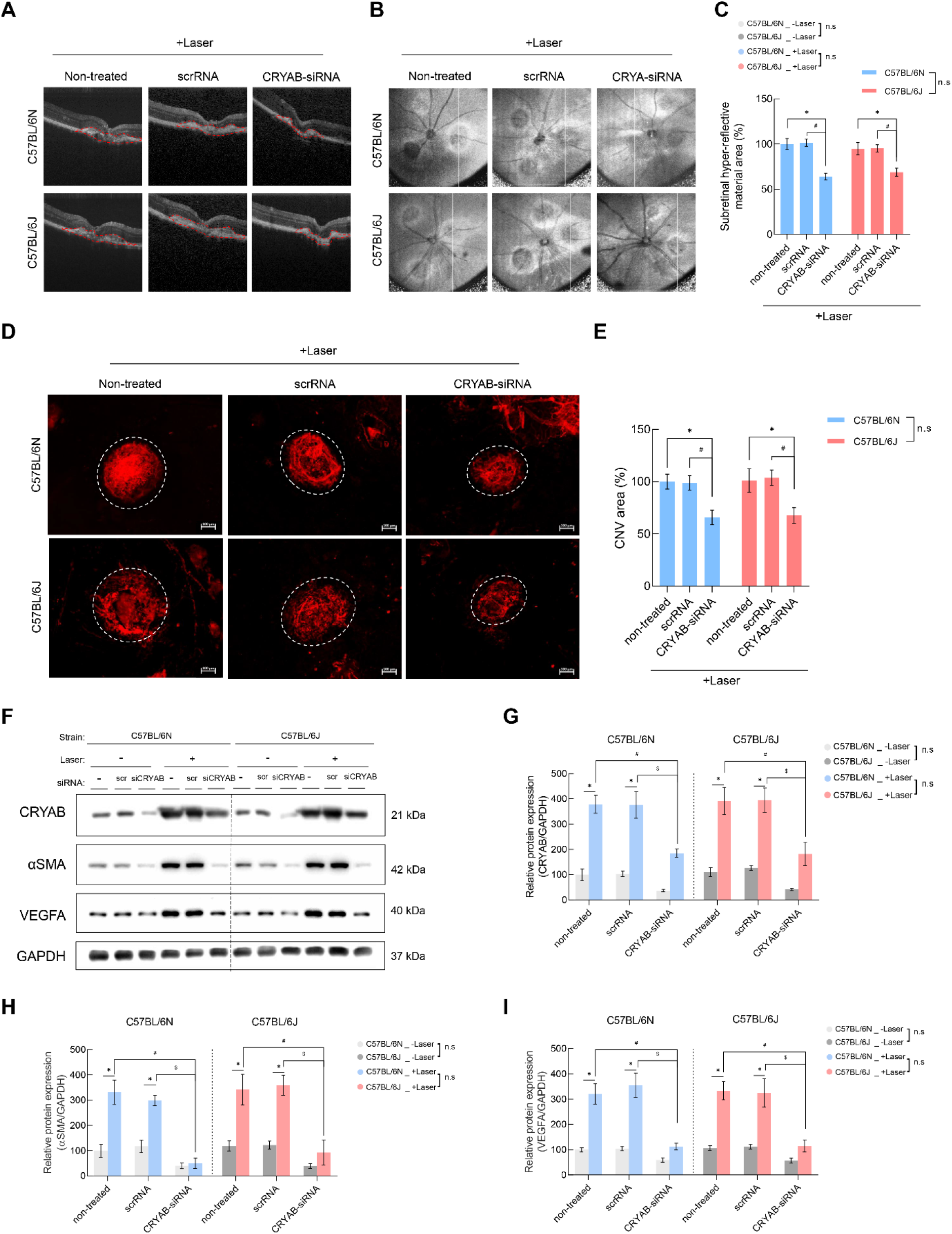
CRYAB knockdown suppresses CNV-associated pathology across C57BL/6 substrains. (A, B) Representative OCT images from laser-induced CNV eyes in C57BL/6N and C57BL/6J mice. Subretinal hyper-reflective lesions are indicated by red dashed outlines. (C) Quantification of the hyper-reflective lesion area corresponding to regions outlined in (A) (n = 18-21). Two-way ANOVA showed a significant effect of siRNA treatment (F(2,117) = 27.93, p < 0.0001), with no significant substrain effect (F(1,117) = 0.3103, p = 0.5785) or siRNA treatment × substrain interaction (F(2,117) = 0.8073, p = 0.4485). Data were analyzed using two-way ANOVA followed by Holm–Šídák’s multiple-comparisons test. Significance is indicated as *p < 0.05 vs. the non-treated group; #p < 0.05 vs. the scrRNA-treated group. (D) Representative images of CNV lesions assessed by IB4-stained RPE/choroid flat mounts in C57BL/6N and C57BL/6J mice following laser injury (with or without CRYAB knockdown). Scale bar, 100 μm. (E) Quantification of CNV area from (D) (n= 10-12). Two-way ANOVA showed a significant effect of siRNA treatment (F(2,64) = 12.85, p < 0.0001), with no significant substrain effect (F(1,64) = 0.1618, p = 0.6888) or siRNA treatment × substrain interaction (F(2,64) = 0.0371, p = 0.9636). Data were analyzed using two-way ANOVA followed by Holm–Šídák’s multiple-comparisons test. Significance is indicated as *p < 0.05 vs. the non-treated group; #p < 0.05 vs. the scrRNA-treated group. (F) Representative immunoblot images showing CRYAB, αSMA, and VEGF expression in laser-induced CNV eyes from C57BL/6N and C57BL/6J mice (with or without CRYAB knockdown). (G–I) Densitometric quantification of (G) CRYAB, (H) αSMA, and (I) VEGFA from immunoblots shown in (F) (n= 6-7). Three-way ANOVA showed significant main effects of siRNA treatment and laser treatment for CRYAB (F(2,62) = 23.72, p < 0.0001; F(1,62) = 156.5, p < 0.0001), αSMA (F(2,62) = 34.98, p < 0.0001; F(1,62) = 68.60, p < 0.0001), and VEGFA (F(2,62) = 30.73, p < 0.0001; F(1,62) = 113.9, p < 0.0001). Significant siRNA treatment × laser interactions were also observed for CRYAB (F(2,62) = 5.540, p = 0.0061), αSMA (F(2,62) = 10.87, p < 0.0001), and VEGFA (F(2,62) = 12.74, p < 0.0001). No significant main effect of substrain or substrain-related interactions was observed for any marker. Significance is indicated as *p < 0.05 vs. non-laser control group; #p < 0.05 vs. −laser and scrRNA treated group; $p < 0.05 vs. +laser and scrRNA treated group.

### 4. Identification of a disease-associated truncated CRYAB species in laser-induced CNV

To investigate whether CRYAB undergoes altered processing during CNV development, we examined CRYAB expression in the laser-induced CNV model (Fig. 4A–E). Laser injury increased VEGFA and αSMA expression and was accompanied by the appearance of an additional truncated CRYAB species. This band was detected predominantly under CNV conditions, indicating that altered CRYAB processing accompanies the response to laser injury.

**Figure 4.**
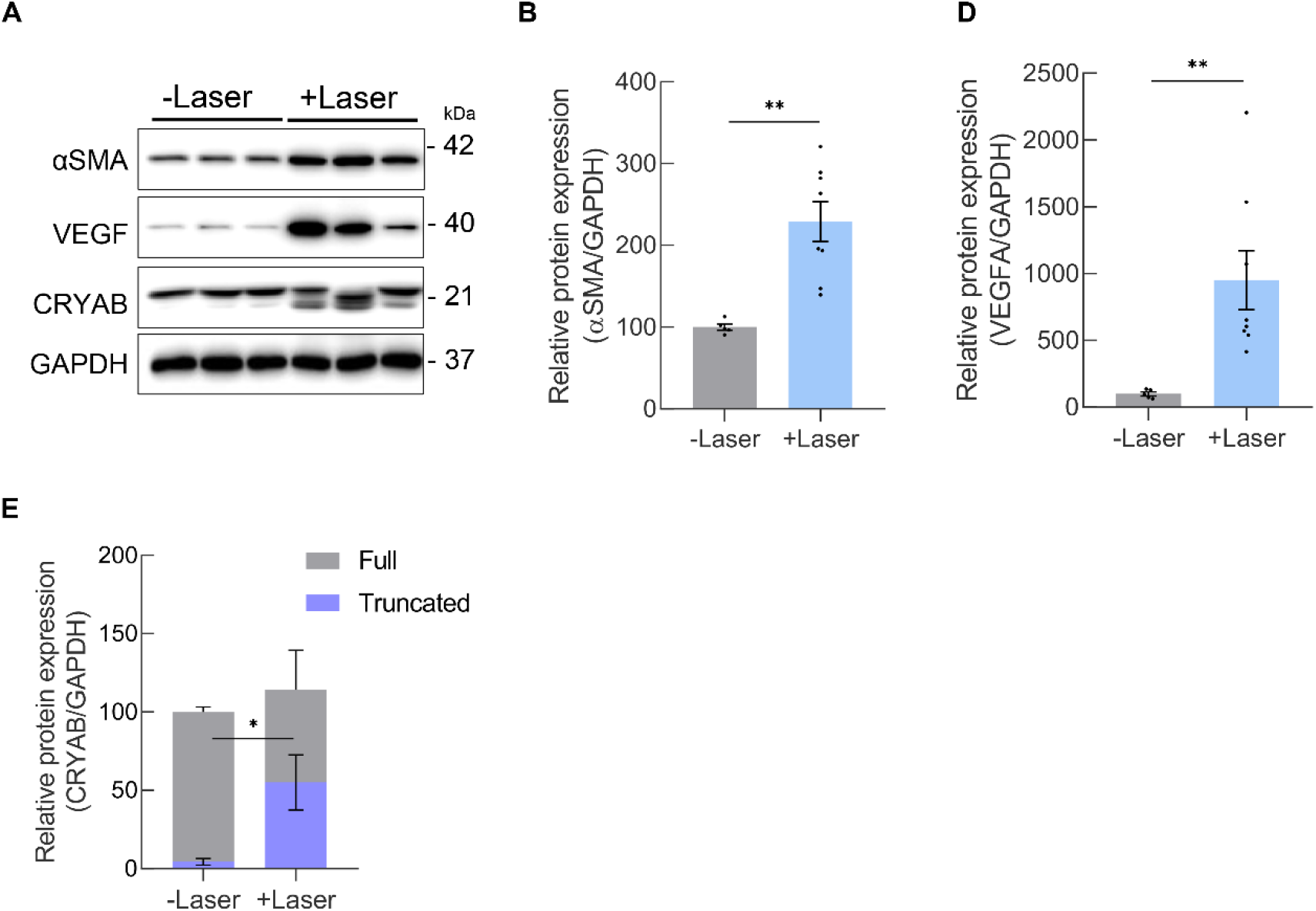
Detection of a Ser45-phosphorylated truncated CRYAB species in experimental CNV. (A) Representative immunoblot images of αSMA, VEGFA, and CRYAB in wild-type (WT) mouse eyes following laser-induced CNV, with GAPDH as a loading control. Quantification of αSMA (B), VEGFA (D), and CRYAB (E) expression is shown (n = 5-8). Data are presented as mean ± SEM and were analyzed using unpaired two-tailed Student’s t-tests. *p < 0.05 and **p < 0.01 vs. non-laser control group.

### 5. Truncated CRYAB bands show reproducible loss of C-terminal peptide coverage

To characterize the molecular features of the truncated CRYAB species, we performed LC–MS/MS analysis on CRYAB-containing gel regions corresponding to full-length and truncated bands isolated from naïve and laser-induced CNV mouse eye lysates (Fig. 5; Fig. S3). In two independent naïve samples, full-length CRYAB was identified with 97% and 96% sequence coverage. Two independent laser-treated samples corresponding to the full-length CRYAB band showed sequence coverage of 97% and 93%, respectively. One laser-treated sample showed a peptide-coverage pattern comparable to that observed in naïve full-length CRYAB, whereas the TIPITR sequence was not identified in the other sample, indicating variability in C-terminal peptide detection among full-length laser-treated samples. The truncated CRYAB bands showed sequence coverage of 88% and 92% in two independent analyses. The C-terminal TIPITR-containing region was not identified in either analysis. In one sample, the entire KQVSGPERTIPITR sequence was not detected, whereas in the other, the K residue together with the TIPITR sequence was not identified. The EFHRK-containing region was consistently not detected across all samples. Overall, the truncated CRYAB bands showed consistent loss of peptide coverage within the distal C-terminal region compared with naïve full-length CRYAB.

**Figure 5.**
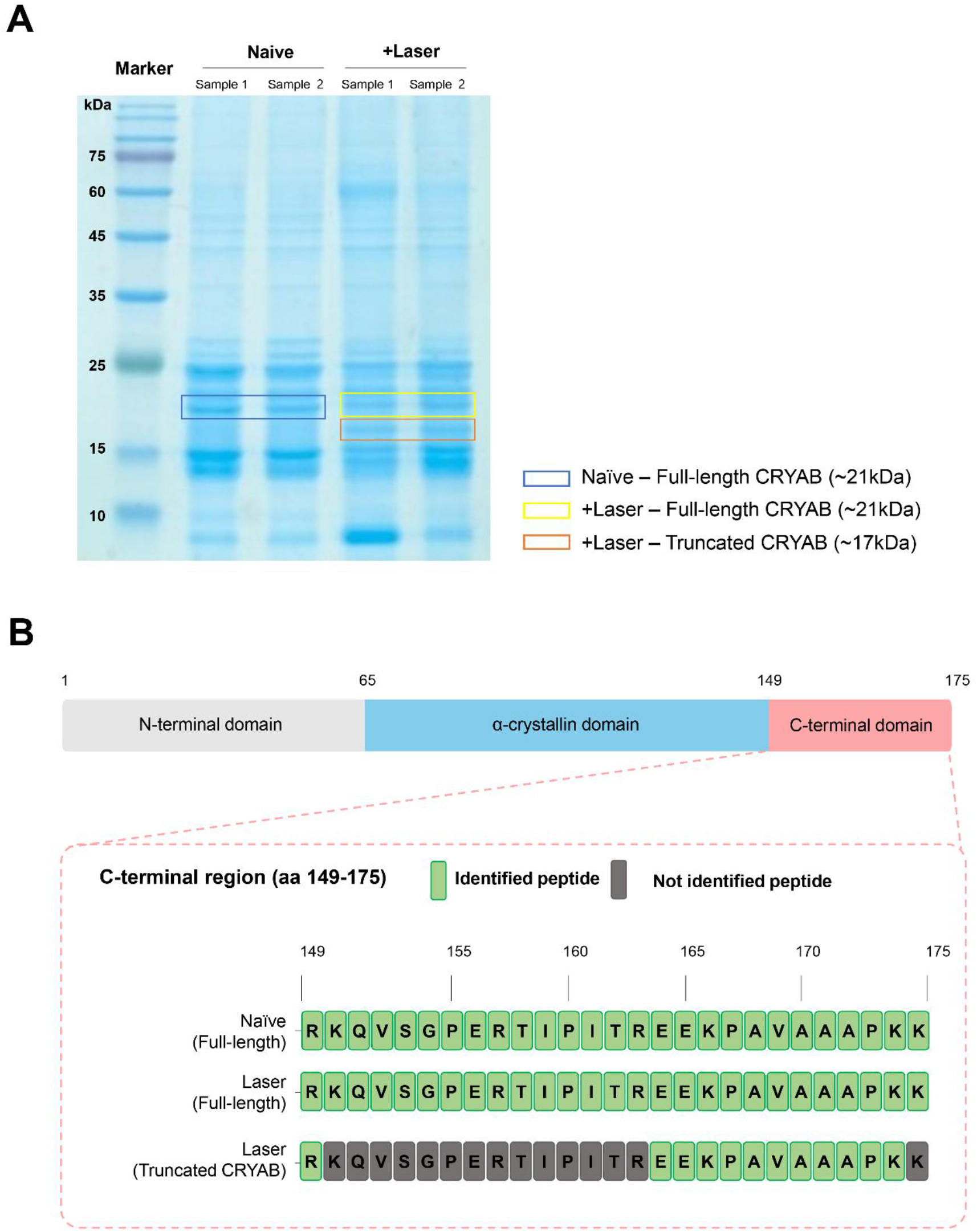
LC–MS/MS peptide mapping of full-length and truncated CRYAB species in mouse eye lysates. (A) Representative Coomassie-stained SDS-PAGE gel showing the regions excised for LC– MS/MS analysis. Gel regions corresponding to full-length CRYAB from naïve eyes (blue), full-length CRYAB from laser-treated eyes (yellow), and truncated CRYAB from laser-treated eyes (orange) were excised for in-gel digestion and subsequent LC–MS/MS analysis. (B) Detailed comparison of LC– MS/MS peptide coverage within the C-terminal region of CRYAB. The C-terminal amino acid sequence is shown for full-length CRYAB from naïve and laser-treated eyes and for truncated CRYAB from laser-treated eyes. Green boxes indicate residues covered by identified peptides, whereas gray boxes indicate residues not identified in the corresponding analysis.

### 6. Differential antibody recognition supports C-terminal alteration of CRYAB in laser-induced CNV

To further characterize the CRYAB species detected in laser-induced CNV, RPE samples from naïve and laser-treated mice were analyzed using two antibodies generated against different CRYAB immunogens (Fig. 6A). Western blotting with an antibody raised against full-length CRYAB detected the expected approximately 21-kDa CRYAB band in both naïve and laser-treated samples (Fig. 6B). Additional bands at approximately 18 and 17 kDa were detected predominantly in laser-treated samples.

**Figure 6.**
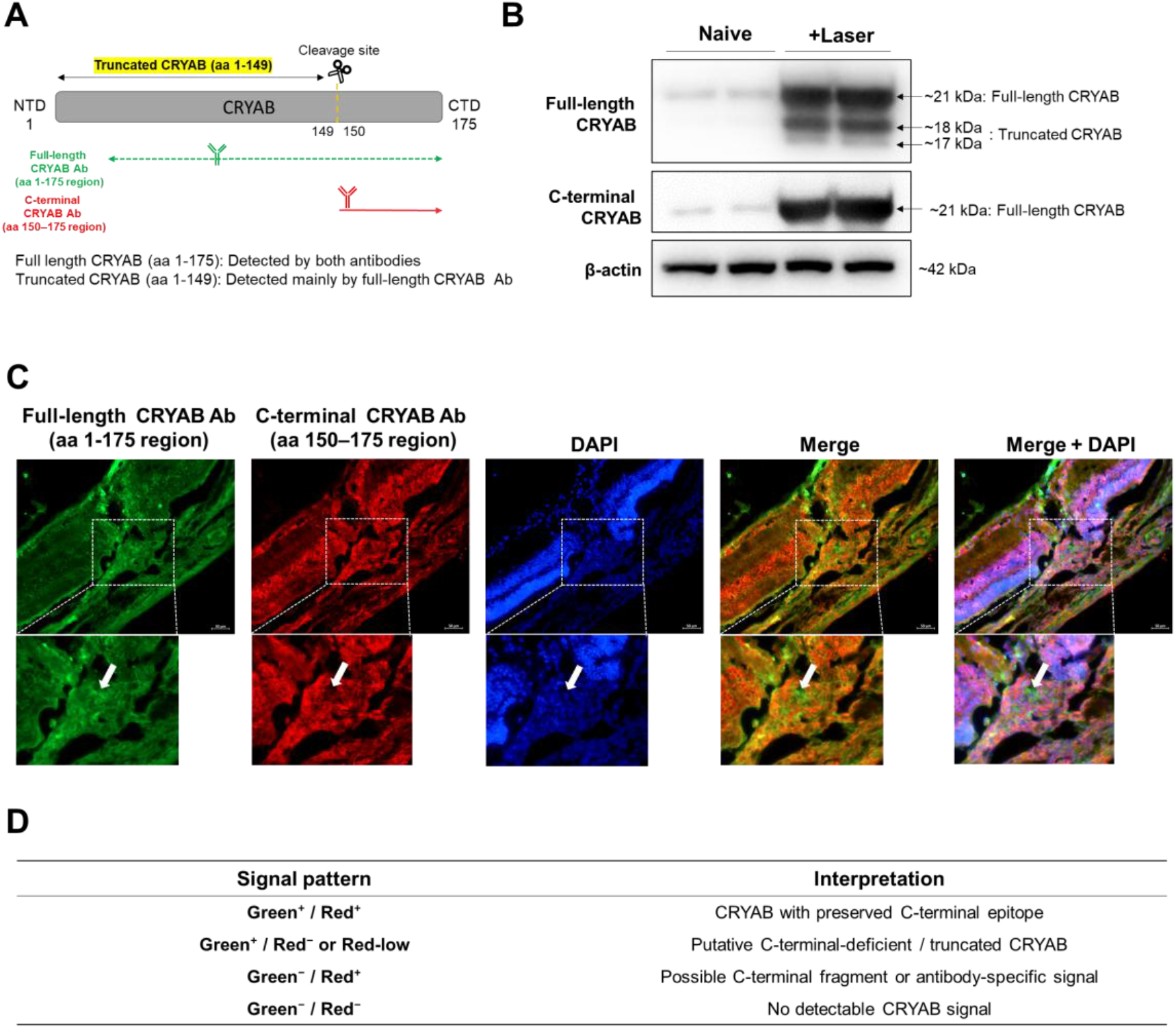
Differential detection of CRYAB species by antibodies raised against distinct immunogens in laser-induced CNV. (A) Western blot analysis of RPE samples from naïve and laser-induced CNV mice using an antibody raised against full-length CRYAB and an antibody generated against a synthetic peptide corresponding to CRYAB amino acids 150 to the C-terminus. The full-length CRYAB antibody detected an approximately 21-kDa band in naïve and laser-treated samples, together with additional approximately 18- and 17-kDa bands in laser-treated samples. The C-terminal antibody detected an approximately 21-kDa band and an additional approximately 19-kDa immunoreactive signal. β-actin was used as a loading control. (B) Representative dual-immunofluorescence images of the retinal lesion area. CRYAB detected using the full-length CRYAB antibody is shown in green, whereas CRYAB detected using the C-terminal antibody is shown in red. Nuclei were counterstained with DAPI (blue). White dashed square indicates green-dominant regions within the lesion area, where the full-length CRYAB antibody signal was retained but the C-terminal antibody signal was weak or absent. (C) The accompanying table summarizes the proposed interpretation of the Western blot and immunofluorescence patterns. The approximately 18- and 17-kDa bands detected only by the native-protein antibody, together with green-dominant/red-low tissue signals, were interpreted as C-terminal-deficient or truncated CRYAB species. Scale bar, 50 μm.

The predicted molecular weight of CRYAB lacking amino acids 150–175 is approximately 17.35 kDa, which is close to the mobility of the approximately 17-kDa band observed in laser-treated samples. In contrast, an antibody generated against a synthetic peptide corresponding to CRYAB amino acids 150 to the C-terminus detected the approximately 21-kDa band and an additional approximately 19-kDa immunoreactive signal but did not detect the approximately 18- and 17-kDa species.

Dual immunofluorescence staining of retinal sections from laser-induced CNV mice showed differential CRYAB immunoreactivity within the lesion area (Fig. 6C,D). Although substantial overlap between the two antibody signals was present, discrete regions showed strong staining with the full-length-CRYAB-immunogen antibody and weak or undetectable staining with the C-terminal-peptide-immunogen antibody.

The differential antibody-recognition pattern, together with the approximate molecular mass of the 17-kDa species, is consistent with loss or alteration of the C-terminal region in a subset of CRYAB molecules under CNV conditions.

### 7. Prss1 is identified among CRYAB-associated proteins under CNV conditions

To investigate proteins associated with CRYAB under CNV conditions, CRYAB immunoprecipitates from laser-induced CNV mouse eyes were subjected to LC– MS/MS-based proteomic analysis (Fig. 7A,B). This analysis identified a subset of CRYAB-associated proteins that were selectively detected in laser-treated samples. Among the proteins identified specifically under CNV conditions was the serine protease Prss1 (Fig. 7C). Co-IP analysis further showed an association between Prss1 and CRYAB-containing complexes in CNV eyes (Fig. 7D), supporting a link between Prss1 and disease-associated CRYAB truncation.

**Figure 7.**
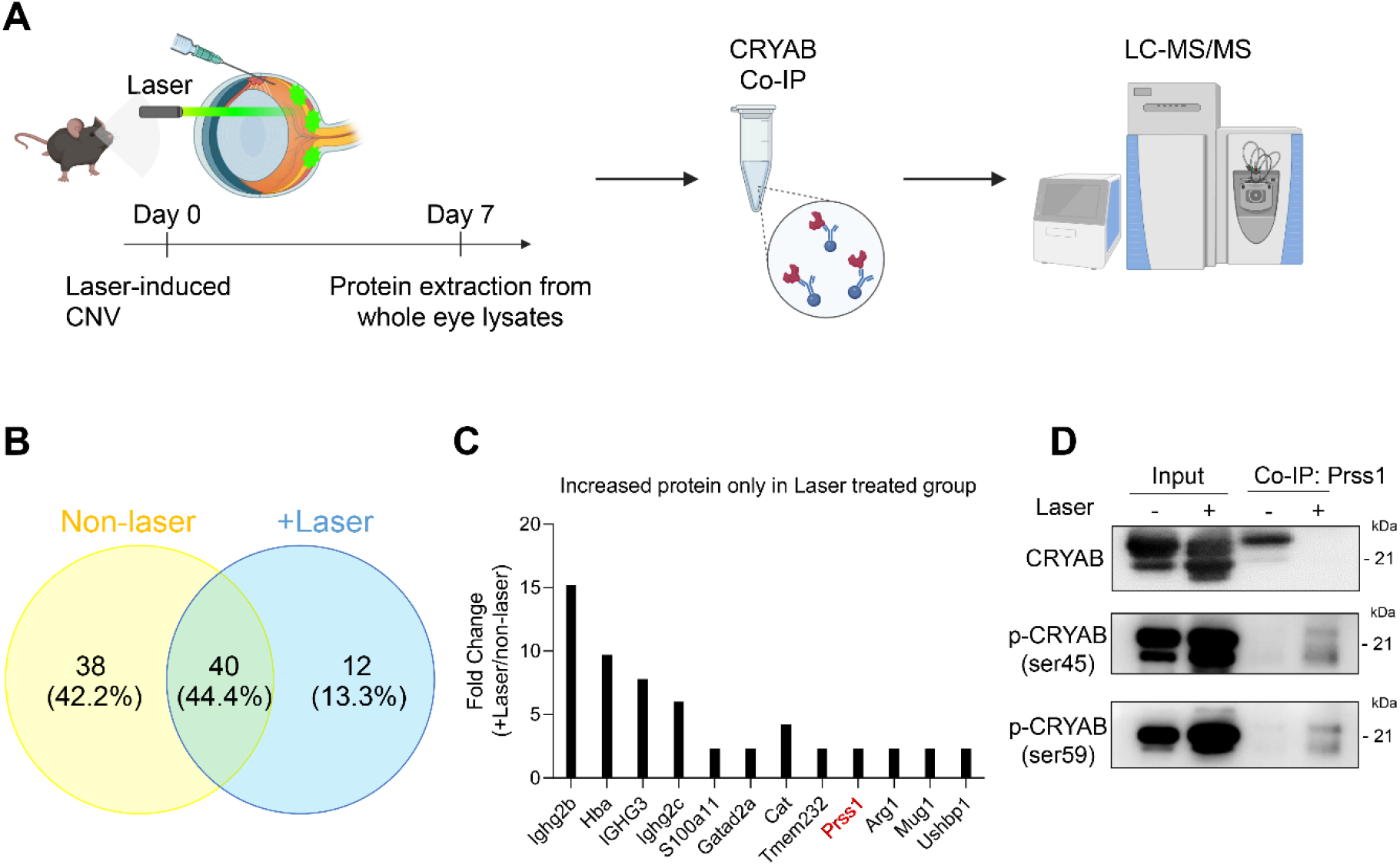
Identification of Prss1 within the CRYAB-associated protease landscape under CNV conditions. (A) Schematic overview of the co-immunoprecipitation (Co-IP) and LC–MS/MS workflow used to identify CRYAB-interacting proteins from non-laser control and laser-induced CNV mouse eyes. (B) Venn diagram summarizing CRYAB-associated proteins identified by LC–MS/MS. Ninety proteins were detected in total, including 40 shared between control and laser-induced samples (green), 38 detected only in control samples (yellow), and 12 detected only in laser-induced CNV samples (blue). (C) Bar graph showing the 12 CRYAB-associated proteins detected specifically in laser-induced CNV samples by LC–MS/MS, including Prss1. (D) Lysates of non-laser and laser-induced mouse eyes were incubated with anti-Prss1 antibodies along with Pierce Protein A/G magnetic beads. Bound proteins were eluted and subjected to western blot with CRYAB and phosphorylated CRYAB (p-CRYAB; Ser45, Ser59).

### 8. Ser45-phosphorylated DTCAMD accumulates with fibrotic activation under inflammation-associated hRPE–THP-1 co-culture conditions

To determine whether the truncated CRYAB species emerges in an inflammation-associated retinal environment, we used a co-culture model consisting of primary human retinal pigment epithelial (hRPE) cells and THP-1-derived macrophages. Truncated Ser45-phosphorylated CRYAB was detected at later time points of co-culture (Fig. 8). Its appearance coincided with a marked increase in αSMA expression, indicating a temporal association between Ser45-phosphorylated DTCAMD accumulation and fibrotic activation.

**Figure 8.**
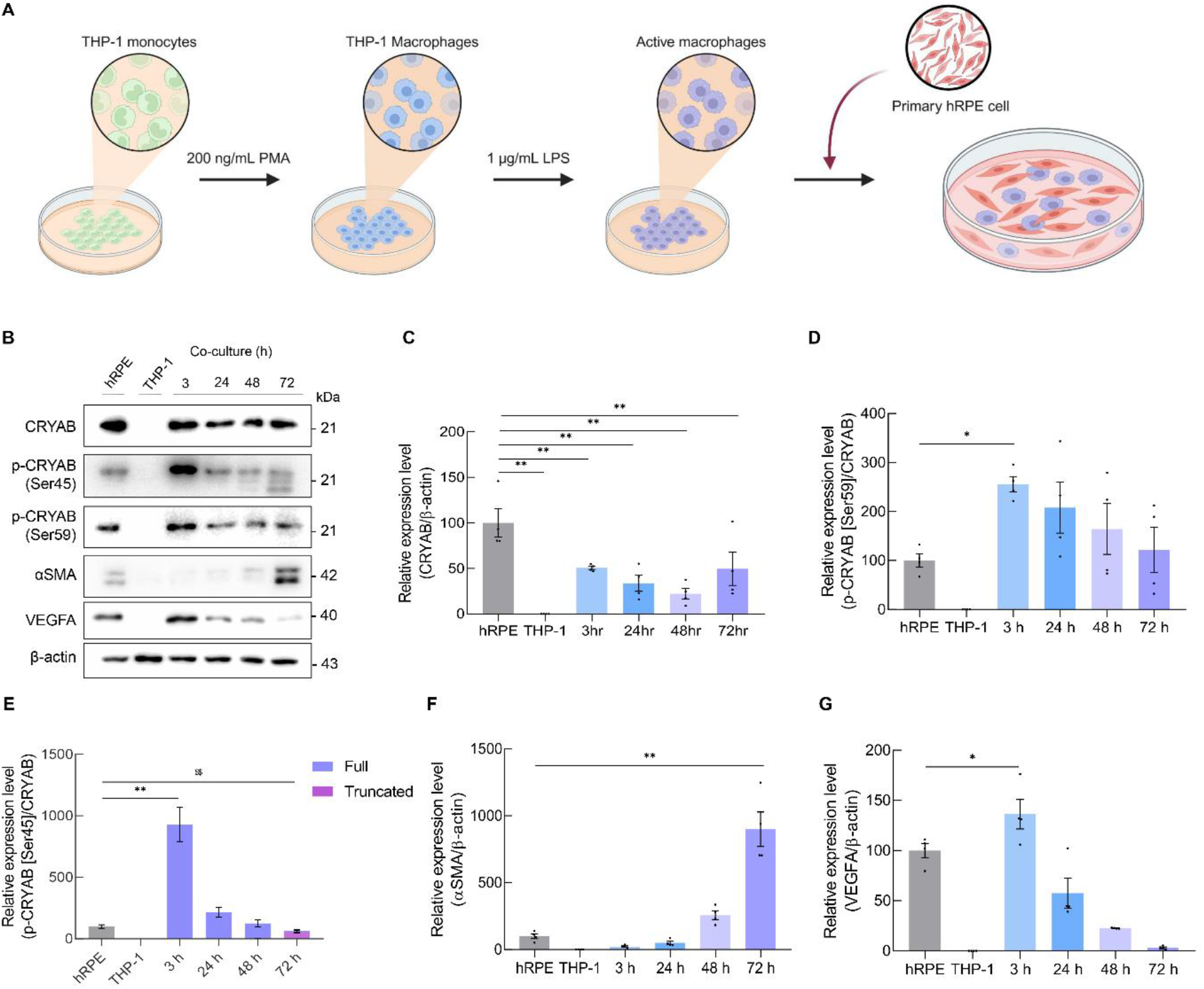
Ser45-phosphorylated truncated CRYAB accumulates with fibrotic activation in hRPE– THP-1 co-culture. (A) Schematic illustration of the primary human RPE and THP-1 co-culture model. The illustration was created in BioRender. Yu, E. (https://BioRender.com/p5mc14x). (B,C) Western blot analysis and quantification of p-CRYAB (Ser45), p-CRYAB (Ser59), total CRYAB, αSMA, and VEGFA in primary hRPE–THP-1 co-cultures at the indicated time points (3, 24, 48, and 72 h). Truncated p-CRYAB (Ser45) was detected at 72 h, as indicated by the appearance of a lower-molecular-weight band. Significant group differences were observed for total CRYAB (F(5,18) = 9.914, p = 0.0001), p-CRYAB (Ser59) (F(5,18) = 6.054, p = 0.0019), full-length p-CRYAB (Ser45) (F(5,18) = 32.78, p < 0.0001), truncated p-CRYAB (Ser45) (F(5,18) = 25.10, p < 0.0001), αSMA (F(5,18) = 39.27, p < 0.0001), and VEGFA (F(5,18) = 37.79, p < 0.0001). Data are presented as mean ± SEM (n = 4) and were analyzed by one-way ANOVA followed by Holm–Šídák’s multiple-comparisons test. *p < 0.05 and **p < 0.01 vs. hRPE (full-length); $$p < 0.01 vs. hRPE (truncated form).

### 9. Pharmacological inhibition of CRYAB processing suppresses DTCAMD accumulation and fibrotic pathology *in vivo*

We next examined whether pharmacological modulation of the CRYAB pathway altered DTCAMD accumulation and CNV-associated pathology *in vivo*. Intravitreal administration of the CRYAB inhibitor T3MG or the serine protease inhibitor AEBSF in laser-induced CNV mice significantly reduced truncated CRYAB and Ser45-phosphorylated DTCAMD signals (Fig. 9A–E).

**Figure 9.**
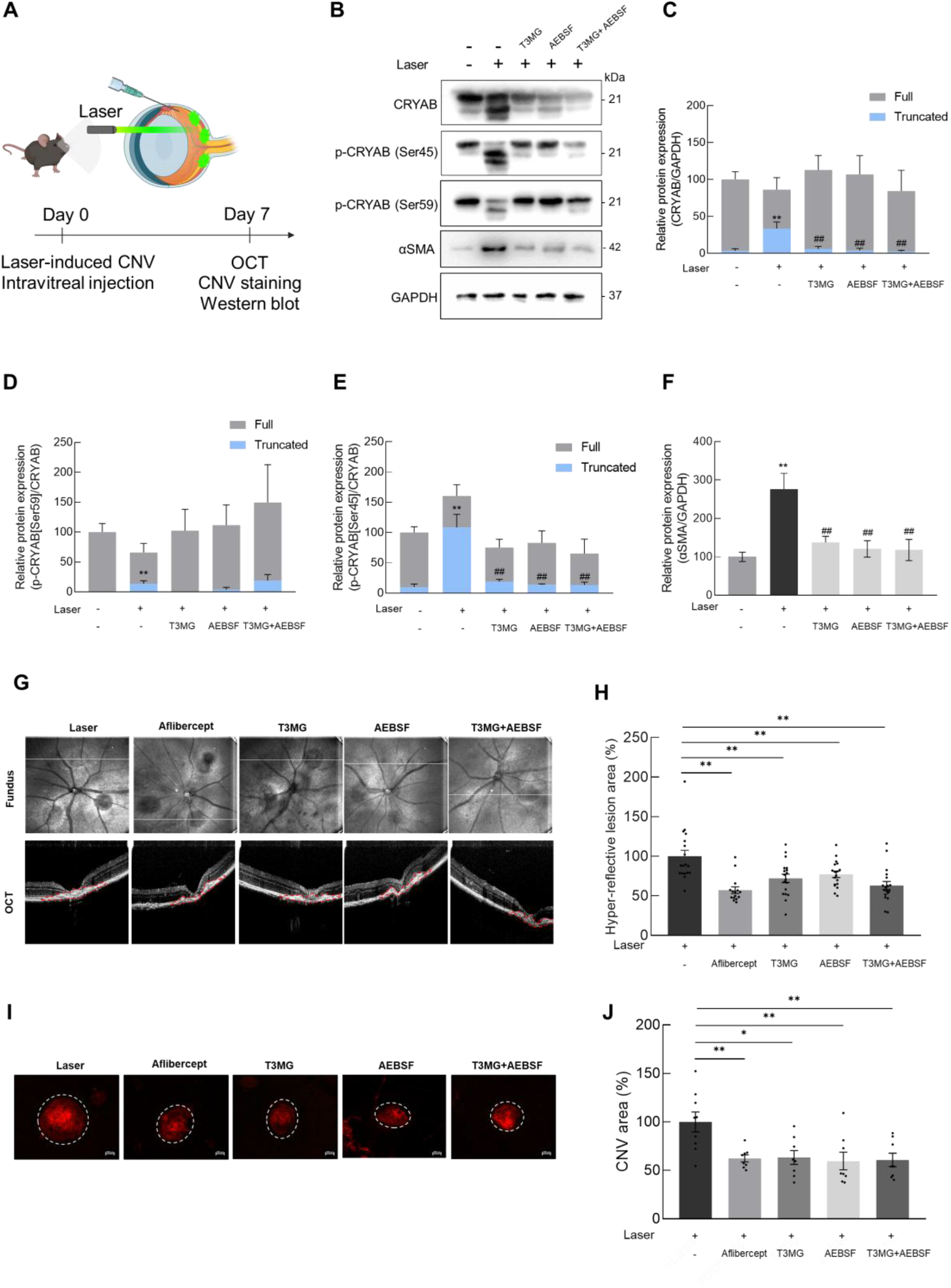
Pharmacological inhibition of CRYAB processing suppresses DTCAMD accumulation and fibrotic pathology in laser-induced CNV mice. (A) Schematic overview of laser-induced CNV induction, intravitreal treatment, and analyses. The illustration was created in BioRender. Yu, E. (https://BioRender.com/p5mc14x). (B) Representative western blot images of CRYAB, phosphorylated CRYAB, and αSMA in eye lysates from each group. Immediately after laser photocoagulation, mice received intravitreal injections of T3MG (25 μg in 1 μL PBS), AEBSF (25 μg in 1 μL PBS), or the combination of T3MG and AEBSF (25 μg in 0.5 μL PBS and 25 μg in 0.5 μL PBS). Equal amounts of protein were analyzed, and GAPDH was used as a loading control. (C–F) Quantification of western blot results for total CRYAB (C), p-CRYAB (Ser59) (D), p-CRYAB (Ser45) (E), and αSMA (F) across the indicated groups. Statistical analysis revealed significant group differences for total CRYAB (F(4,22) = 7.434, p = 0.0006), p-CRYAB (Ser59) (Kruskal–Wallis H = 12.46, p = 0.0142), p-CRYAB (Ser45) (F(4,22) = 15.17, p < 0.0001), and αSMA (F(4,22) = 7.714, p = 0.0005). Data are presented as mean ± SEM and were analyzed using one-way ANOVA or the Kruskal–Wallis test, as appropriate (*p < 0.05 and **p < 0.01 vs. the non-laser control group; #p < 0.05 and ##p < 0.01 vs. the laser-only group). (G) Representative OCT images showing CNV-associated subretinal hyper-reflective lesions in each group. Red dotted lines denote the lesion areas. (H) Quantification of hyper-reflective lesion area from OCT images in (G) (n = 15-18). Data are presented as mean ± SEM and were analyzed by one-way ANOVA followed by Holm–Šídák’s multiple-comparisons test (F(4,82) = 8.886, p < 0.0001). (I) Representative images of CNV lesions in IB4-stained choroidal flat mounts. White dotted lines denote the lesion areas. Scale bar, 100 μm. (J) Quantification of CNV lesion area from IB4-stained choroidal flat mounts in (I) (n = 8-9). Data are presented as mean ± SEM and were analyzed by one-way ANOVA followed by Holm– Šídák’s multiple-comparisons test (F(4,36) = 5.111, p = 0.0023).

These changes were accompanied by decreased αSMA expression, CNV area, and subretinal hyper-reflective lesion area (Fig. 9F–J). Both pharmacological interventions therefore reduced DTCAMD-associated signals together with angiogenic and fibrotic readouts in the laser-induced CNV model. The experiments establish an association between pharmacological suppression of the CRYAB/protease axis and reduced pathological responses, while the precise contribution of DTCAMD processing to each outcome requires further mechanistic validation.

## Discussion

CRYAB has been implicated in epithelial–mesenchymal transition (EMT), a central component of fibrotic remodeling that encompasses changes in cell phenotype, migration, and extracellular matrix production^25–28^. Consistent with this, CRYAB overexpression has been reported to promote EMT-like responses, whereas CRYAB suppression favors mesenchymal-to-epithelial transition and reduces subretinal fibrosis in experimental models^27^. Elevated CRYAB levels have also been described across multiple retinal disorders, including rhegmatogenous retinal detachment, proliferative diabetic retinopathy, and proliferative vitreoretinopathy^29–31^. Despite these observations, how CRYAB shifts from a predominantly cytoprotective chaperone toward a disease-associated pro-fibrotic role in the retina remains unclear.

Here, we show that CRYAB is increased in the aqueous humor of patients with nAMD and is associated with early anatomical response to anti-VEGF therapy. In our cohort, aqueous humor CRYAB showed a higher ROC AUC than the OCT-derived parameters examined, suggesting that CRYAB reflects aspects of disease activity not fully captured by structural imaging alone. These findings extend the clinical relevance of CRYAB beyond its increased abundance and point to its potential value as a molecular indicator of treatment response.

The functional contribution of CRYAB to CNV-associated pathology was supported by complementary genetic approaches. CRYAB-deficient mice exhibited reduced CNV leakage and hyper-reflective lesion formation, together with decreased VEGFA and αSMA expression, compared with WT mice, supporting a functional role for CRYAB in both angiogenic and fibrotic responses. This finding was reinforced by siRNA-mediated CRYAB knockdown, which reduced VEGFA and αSMA expression under laser-induced CNV conditions in both C57BL/6N and C57BL/6J mice. The absence of significant substrain effects or substrain-dependent interactions further indicates that the effects of CRYAB suppression are reproducible across C57BL/6 backgrounds. These genetic data establish CRYAB as an active component of CNV-associated pathology and provided the basis for examining whether disease-associated molecular forms of CRYAB emerge during lesion development.

A major observation was the appearance of a truncated CRYAB species under disease conditions. This species was detected in experimental CNV, where it was associated with Ser45 phosphorylation and fibrotic marker expression. We refer to this disease-associated species as DTCAMD. Its preferential appearance under CNV conditions suggests that pathological CRYAB activity may involve qualitative changes in protein processing in addition to increased expression.

The molecular features of this truncated CRYAB species were further examined by LC–MS/MS peptide mapping. Compared with full-length CRYAB, the truncated bands showed reproducible loss of peptide coverage within the distal C-terminal region. In both independent analyses of the truncated species, K150 and the TIPITR-containing region were not identified. This reproducible loss of distal C-terminal peptide coverage was consistent with C-terminal alteration of CRYAB. Complementary antibody-based analyses provided additional evidence for this alteration. In western blotting, an antibody raised against full-length CRYAB detected additional bands of approximately 18 and 17 kDa in laser-induced CNV samples, whereas an antibody generated against CRYAB peptide spanning amino acid 150 to the C-terminus did not detect these species. The approximately 17-kDa band also closely matched the predicted molecular mass of CRYAB lacking residues 150–175. Dual immunofluorescence revealed lesion-associated regions in which signal detected by the full-length-CRYAB-immunogen antibody was retained, whereas signal obtained with the C-terminal-peptide-immunogen antibody was weak or absent. Collectively, these results indicate accumulation of a C-terminally altered CRYAB species under CNV conditions.

C-terminal alteration could also affect the structural organization of CRYAB. The C-terminal domain (CTD) of CRYAB contains a conserved IXI/V motif that interacts with the β4–β8 hydrophobic groove within the α-crystallin domain (ACD), thereby contributing to oligomeric organization and regulation of intermolecular interactions^32^. Loss of the distal C-terminal segment would remove this motif and could alter accessibility of the β4–β8 groove. Such a model is consistent with findings by Hafizi et al., who showed that structural perturbation of CRYAB, including the D109A mutation, disrupts stabilizing interactions and increases hydrophobic surface exposure and aggregation^18^. In this context, truncation of the C-terminal region provides a potential mechanistic basis by which disease-associated processing could modify the interaction profile of CRYAB and favor non-canonical protein associations.

Our findings further place DTCAMD formation within a protease-associated pathological environment. Proteolytic processing has been implicated in ocular disease mechanisms; for example, HTRA1-mediated cleavage of thrombospondin-1 has been reported to compromise anti-angiogenic activity^33^. Among CRYAB-associated proteins identified under CNV conditions, Prss1 was selectively detected in laser-treated samples and its association with CRYAB-containing complexes was supported by Co-IP. Prss1 is a trypsin-like serine protease, and the C-terminal loss inferred from peptide mapping occurs within a region containing Arg149 and Lys150, residues compatible with trypsin-like substrate specificity. The reduction of DTCAMD-associated signals following AEBSF treatment further links serine protease activity to CRYAB truncation. These observations identify Prss1 as a plausible candidate protease involved in DTCAMD formation, although direct biochemical studies will be required to establish the enzyme–substrate relationship and define the precise truncation site.

DTCAMD accumulation was also closely associated with fibrotic activation. In the hRPE–THP-1 co-culture model, a truncated Ser45-phosphorylated CRYAB species became evident at later time points together with increased αSMA expression, and a similar pattern was observed *in vivo*. These data link Ser45-phosphorylated DTCAMD with a fibrotic cellular state but do not establish that Ser45 phosphorylation drives truncation or αSMA induction. Ser45 phosphorylation is regulated by the p44/42 MAPK pathway,^34–36^, which participates in cellular stress responses and extracellular matrix remodeling^37,38^, whereas Ser59 phosphorylation has been linked more strongly to inflammatory signaling and cell-survival pathways^34,36,39,40^. The preferential appearance of the truncated species in association with Ser45 phosphorylation therefore raises the possibility that phosphorylation state influences CRYAB processing or its interaction profile during fibrotic remodeling.

The genetic findings were complemented by pharmacological modulation of the CRYAB pathway. T3MG^23,41^ and the serine protease inhibitor AEBSF reduced truncated CRYAB/DTCAMD-associated signals, αSMA expression, CNV area, and subretinal hyper-reflective lesions in the laser-induced CNV model. Co-administration also reduced these pathological readouts, supporting the concept that CRYAB activity and protease-associated processing represent interconnected components of the same pathological axis. Although T3MG and AEBSF are experimental tools rather than clinical candidates, their effects provide pharmacological support for the genetic data and demonstrate that modulation of the CRYAB–protease axis can suppress both DTCAMD accumulation and CNV-associated angiogenic and fibrotic responses.

Several limitations should be considered. The clinical analysis was based on a relatively small cohort, and the association between aqueous humor CRYAB and treatment response requires validation in larger prospective populations. Although LC–MS/MS peptide mapping and differential antibody recognition support C-terminal alteration, neither approach defines the exact truncation site. Likewise, the association of Prss1 with CRYAB does not establish Prss1 as the protease responsible for CRYAB truncation. The laser-induced CNV model also represents an acute injury response and does not fully reproduce the chronic evolution of human nAMD and mature subretinal fibrosis. Direct biochemical studies of CRYAB truncation, structural characterization of DTCAMD, and longer-term fibrosis models will be important for resolving these questions.

In conclusion, our study identifies CRYAB as a functional contributor to CNV-associated angiogenic and fibrotic pathology and describes a disease-associated, Ser45-phosphorylated truncated CRYAB species that accumulates under fibrotic conditions. Genetic deletion and knockdown of CRYAB suppressed VEGFA and αSMA responses, while peptide mapping and differential antibody recognition support C-terminal alteration of CRYAB. The association with Prss1 and the sensitivity of DTCAMD accumulation to serine protease inhibition further implicate proteolytic processing in the generation of this disease-associated CRYAB species. These findings define a CRYAB processing axis linking protein modification to CNV-associated fibrotic remodeling and provide a mechanistic basis for targeting pathological CRYAB processing in nAMD.

## Supporting information

BioRender Publication License

## Authorship contribution statement

**E.Y.**: Investigation; Formal analysis; Data curation; Methodology; Validation; Visualization; Writing – original draft. **S.M.G.**: Formal analysis; Data curation. **J.J.**: Investigation; Formal analysis; Methodology; Data curation. **J.K.M.**: Resources; Methodology; Data curation; Project administration; Funding acquisition; Writing – review & editing. **J.Y.**: Conceptualization; Resources; Methodology; Project administration; Writing – review & editing; Supervision; Funding acquisition.

