## Supplemental figures and legends for "DTCAMD, a disease-associated truncated form of CRYAB in neovascular age-related macular degeneration: a potential driver of subretinal fibrosis"

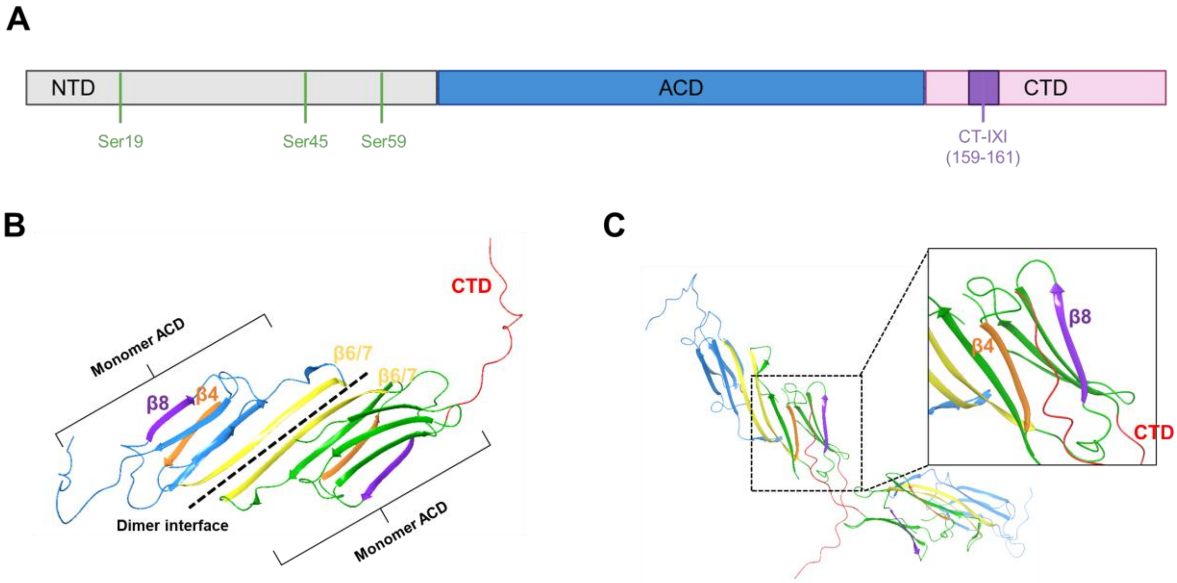


**Fig. S 1. Structural organization of CRYAB and CTD engagement of the ACD β4–β8 groove.**
(A) Linear domain map highlighting the N-terminal region (NTR), α-crystallin domain (ACD), and C-terminal domain (CTD) containing the conserved IXI/V motif (Ile159–Pro160–Ile161). (B) Structure of the CRYAB ACD dimer. The dimer interface is formed by the β6/β7 strands of each monomer (β6/β7 shown in yellow). (C) Structural overview of CRYAB dimer–dimer arrangement illustrating CTD insertion into the ACD β4–β8 groove. The CTD (red) from a neighboring subunit occupies the groove formed between β4 (orange) and β8 (purple).

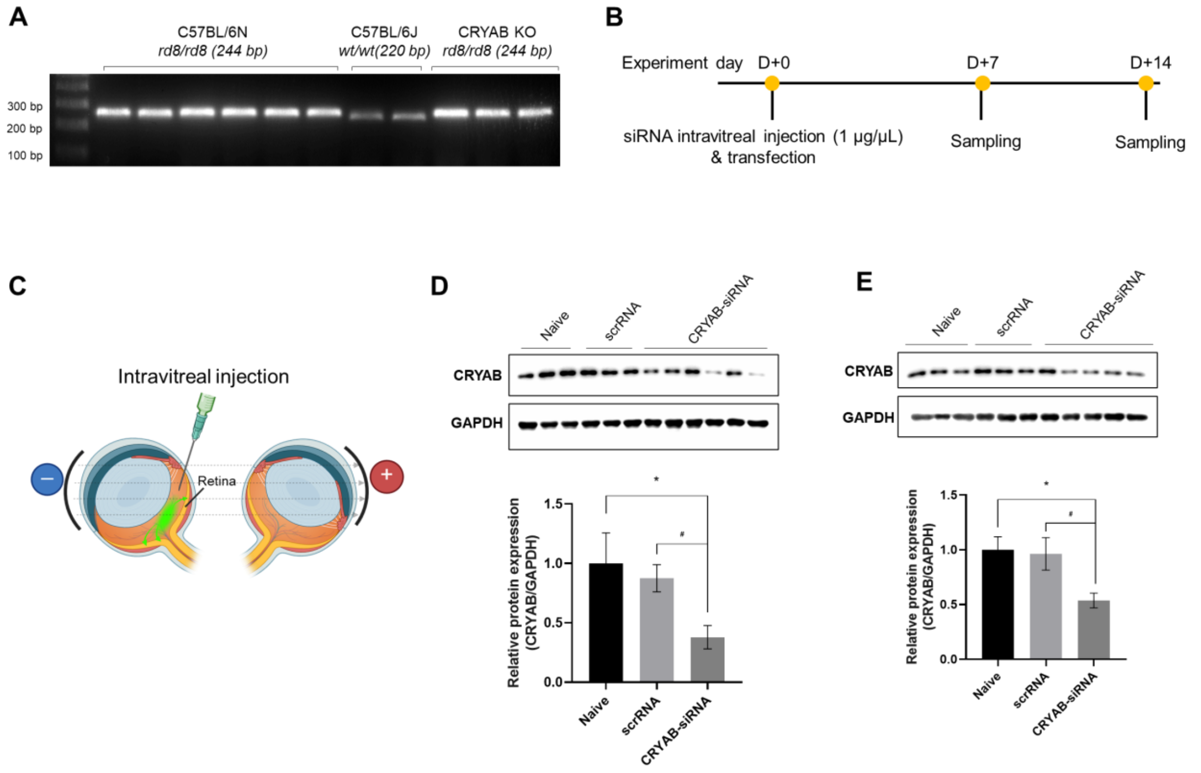


**Fig. S 2.** ***In vivo* electroporation workflow and validation of CRYAB knockdown efficiency.** (A) Representative PCR genotyping of C57BL/6 substrains and CRYAB knockout mice, showing the *Crb1*$^{rd8}$ allele in C57BL/6N mice and C57BL/6N-background CRYAB KO mice (rd8/rd8, 244 bp) and the wild-type allele in C57BL/6J mice (wt/wt, 220 bp). (B) Experimental timeline for intravitreal siRNA delivery followed by in vivo electroporation and tissue collection at 7 and 14 days post-transfection. (C) Schematic illustration of electrode placement during electroporation to facilitate siRNA delivery to the RPE. The cathode (negative electrode) was positioned on the injected (left) eye, and the anode (positive electrode) was placed on the contralateral (right) eye. (D) Representative immunoblotting results showing CRYAB expression at day 7 after electroporation in eyes receiving CRYAB-specific siRNA or scrambled control siRNA. GAPDH was used as a loading control (n = 3-6). (E) Representative immunoblotting results showing CRYAB expression at day 14 after electroporation in eyes receiving CRYAB-specific siRNA or scrambled control siRNA. GAPDH was used as a loading control (n = 3-6). Data are presented as mean ± SEM and analyzed by one-way ANOVA (*p < 0.05 vs. the non-treated group; #p < 0.05 versus the scrRNA treated group).

## A

### Naïve full-length CRYAB

**Sample 1**

**Protein sequence coverage: 97%**

Matched peptides shown in ***bold red***.

```
  1 MDIAIHHPWI RRPFFPFHSP SRLFDQFFGE HLLESDLFST ATSLSPFYLR
 51 PPSFLRAPSW IDTGLSEMRL EKDRFSVNLD VKHFSPEELK VKVLGDVIEV
101 HGKHEERQDE HGFISREFHR KYRIPADVDP LTITSSLSSD GVLTVNGPRK
151 QVSGPERTIP ITREEKPAVA AAPKK
```

Unformatted sequence string: 175 residues (for pasting into other applications).

**Sample 2**

**Protein sequence coverage: 96%**

Matched peptides shown in ***bold red***.

```
  1 MDIAIHHPWI RRPFFPFHSP SRLFDQFFGE HLLESDLFST ATSLSPFYLR
 51 PPSFLRAPSW IDTGLSEMRL EKDRFSVNLD VKHFSPEELK VKVLGDVIEV
101 HGKHEERQDE HGFISREFHR KYRIPADVDP LTITSSLSSD GVLTVNGPRK
151 QVSGPERTIP ITREEKPAVA AAPKK
```

Unformatted sequence string: 175 residues (for pasting into other applications).

## B

### Laser-treated full-length CRYAB

**Sample 1**

**Protein sequence coverage: 93%**

Matched peptides shown in ***bold red***.

```
  1 MDIAIHHPWI RRPFFPFHSP SRLFDQFFGE HLLESDLFST ATSLSPFYLR
 51 PPSFLRAPSW IDTGLSEMRL EKDRFSVNLD VKHFSPEELK VKVLGDVIEV
101 HGKHEERQDE HGFISREFHR KYRIPADVDP LTITSSLSSD GVLTVNGPRK
151 QVSGPERTIP ITREEKPAVA AAPKK
```

Unformatted sequence string: 175 residues (for pasting into other applications).

**Sample 2**

**Protein sequence coverage: 97%**

Matched peptides shown in ***bold red***.

```
  1 MDIAIHHPWI RRPFFPFHSP SRLFDQFFGE HLLESDLFST ATSLSPFYLR
 51 PPSFLRAPSW IDTGLSEMRL EKDRFSVNLD VKHFSPEELK VKVLGDVIEV
101 HGKHEERQDE HGFISREFHR KYRIPADVDP LTITSSLSSD GVLTVNGPRK
151 QVSGPERTIP ITREEKPAVA AAPKK
```

Unformatted sequence string: 175 residues (for pasting into other applications).

## C

### Laser-treated truncated CRYAB

**Sample 1**

**Protein sequence coverage: 88%**

Matched peptides shown in ***bold red***.

```
  1 MDIAIHHPWI RRPFFPFHSP SRLFDQFFGE HLLESDLFST ATSLSPFYLR
 51 PPSFLRAPSW IDTGLSEMRL EKDRFSVNLD VKHFSPEELK VKVLGDVIEV
101 HGKHEERQDE HGFISREFHR KYRIPADVDP LTITSSLSSD GVLTVNGPRK
151 QVSGPERTIP ITREEKPAVA AAPKK
```

Unformatted sequence string: 175 residues (for pasting into other applications).

**Sample 2**

**Protein sequence coverage: 92%**

Matched peptides shown in ***bold red***.

```
  1 MDIAIHHPWI RRPFFPFHSP SRLFDQFFGE HLLESDLFST ATSLSPFYLR
 51 PPSFLRAPSW IDTGLSEMRL EKDRFSVNLD VKHFSPEELK VKVLGDVIEV
101 HGKHEERQDE HGFISREFHR KYRIPADVDP LTITSSLSSD GVLTVNGPRK
151 QVSGPERTIP ITREEKPAVA AAPKK
```

Unformatted sequence string: 175 residues (for pasting into other applications).

**Fig. S 3. Mascot peptide-mapping results for full-length and putative truncated CRYAB.**
Mascot protein views are shown for two independent samples of naïve full-length CRYAB, laser-treated full-length CRYAB, and laser-treated putative truncated CRYAB. Sequence coverage was 97% and 96% in naïve full-length samples (A), 97% and 93% in laser-treated full-length samples (B), and 88% and 92% in truncated CRYAB samples (C). Matched peptides are indicated in red in the Mascot sequence display. The TIPITR-containing C-terminal region was not identified in either putative truncated CRYAB sample, whereas the EFHRK-containing region was not detected in any analyzed sample.
