## Supplementary material for "DTCAMD, a disease-associated truncated form of CRYAB in neovascular age-related macular degeneration: a potential driver of subretinal fibrosis": BioRender Publication License: 4 - Publication License Oct-02-2026.pdf

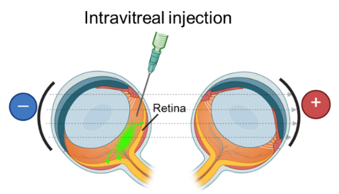


*For any questions regarding this document, or other questions about publishing with BioRender, please refer to our BioRender Publication Guide, or contact BioRender Support at.*
